# Navigating uncertainty: reward location variability induces reorganization of hippocampal spatial representations

**DOI:** 10.1101/2025.01.06.631465

**Authors:** Charline Tessereau, Feng Xuan, Zehua Chen, Jack R. Mellor, Peter Dayan, Daniel Dombeck

## Abstract

Navigating uncertainty is crucial for survival, with the location and availability of reward varying in different and unsignalled ways. Hippocampal place cell populations over-represent salient locations in an animal’s environment, including those associated with rewards; however, how the spatial uncertainties impact the cognitive map is unclear. We report a virtual spatial navigation task designed to test the impact of different levels and types of uncertainty about reward on place cell populations. When the reward location changed on a trial-by-trial basis, inducing expected uncertainty, a greater proportion of place cells followed along, and the reward and the track end became anchors of a warped spatial metric. When the reward location then unexpectedly moved, the fraction of reward place cells that followed was greater when starting from a state of expected, compared to low, uncertainty. Overall, we show that different forms of potentially interacting uncertainty generate remapping in parallel, task-relevant, reference frames.

## Introduction

Animals, including humans, thrive according to their ability to adapt to tasks, situations, and environments which vary in their regularity and associated uncertainties. For instance, while driving, minor unpredictable delays are common, would not prompt a route change and may even be unnoticed. They can thus be considered a form of *expected uncertainty* (often associated with aleatoric uncertainty or risk; Hüllermeier and Waegeman, 2021), in which precise outcomes are not fully forseeable. However, a traffic jam can seem very surprising for someone used to a clear commute, a form of *unexpected uncertainty* (Soltani and Izquierdo, 2019; Yu and Dayan, 2005). This can indicate a significant contextual change that might necessitate significant adaptation, for instance, the need to use another route. Importantly, the threshold to consider an outcome as unexpected differs depending on expected uncertainty, for example, sporadic traffic jams might be customary to someone living in a busy capital, but prompting unexpected uncertainty in a small countryside town. For the brain to process and interpret these interacting forms of uncertainty is critical for adaptive behavior.

Most research on neural correlates of uncertainties has concentrated on aspects of decision-making, related to rewards and punishments (Behrens et al, 2007; Cohen et al, 2015; Dayan, 2012; Hsu et al, 2005; McGuire et al, 2014; Nassar et al, 2019; Preuschoff et al, 2011; Soltani and Izquierdo, 2019; Yu and Dayan, 2005). By contrast, it has rarely been applied to spatial contexts such as the location-specific traffic example above. In particular, the concept of uncertainty has not previously been applied to the description and understanding of spatial representations in the hippocampus and related structures, such as the well-studied place cells (Bast et al, 2009; Best et al, 2001; Burgess et al, 1995; Dombeck et al, 2010; Kleinknecht et al, 2012; Morris et al, 1990; Moser et al, 2008; Muller, 1996; O’Keefe and Dostrovsky, 1971; Radvansky et al, 2021; Sosa and Giocomo, 2021; Tessereau et al, 2021)(O’Keefe and Dostrovsky, 1971), even though many previous results might fit into such a general framework. For example, whether the hippocampal place cell population (i.e. the cognitive map) changes gradually or suddenly during a progressive change to the features (e.g. shape) of an animal’s environment depends on the amount of experience the animal has had with the intermediate features (Leutgeb et al, 2005a; Plitt and Giocomo, 2021; Wills et al, 2005): the more experience, the more expected uncertainty and the more gradually the place cell population changes; the less experience, the less expected uncertainty and the more suddenly the place cell population changes. Though previous hippocampal research did not explicitly describe results in terms of uncertainty, insights for understanding how place cell populations might map environments with different levels of uncertainty can still be deduced.

In the case of expected uncertainty, for example, varying the spatial environment on a trial-by-trial basis (i.e., expected uncertainty in the spatial reference frame) caused hippocampal activity to reflect the statistics of the episodic environment (Plitt and Giocomo, 2021). Perhaps similarly, switching a stable reward location by block (e.g. expected uncertainty in reward location on a timescale of tens of minutes timescale) induces the progressive recruitment of reward-centred place cells (Gauthier and Tank, 2018; Issa et al, 2024; Sosa et al, 2023). However, reward foraging behaviors in nature often involve rapid, non-random, changes in reward locations in a stable spatial environment, a condition of expected uncertainty that has not been explored in prior studies. Therefore, it is unclear how the hippocampal place cell population encodes expected uncertainty in reward location on a trial by trial basis, independent of changes to the spatial reference frame.

In the case of unexpected uncertainty, numerous prior studies provide insights into how place cell populations change their encoding when animals are exposed to large, unexpected changes to their environments. This is typically induced by switching the animals to a novel arena or track which bears little resemblance to previously experienced spaces. These manipulations often result in a phenomenon known as remapping (Anderson and Jeffery, 2003; Bostock et al, 1991; Kentros et al, 1998; Leutgeb et al, 2005b; Muller and Kubie, 1987; Sanders et al, 2020), where place fields change their activity patterns between the two environments (Frank et al, 2004; Hill, 1978; Michon et al, 2021; Sheffield et al, 2017; Wills et al, 2005). While such ”remapping” experiments are typically performed by changing aspects of space, prior studies have not looked at unexpected uncertainty in reward location, independent of changes to the spatial reference frame, without prior experience for such a move. Furthermore, in prior ”remapping” experiments, the level of uncertainty between the familiar and novel experiences has not been systematically varied. Thus, not only is it not clear how changes to hippocampal representations in the light of unexpected and expected uncertainty compare, but it is also unknown whether the encoding changes to place cells that are induced by unexpected uncertainty depend on the initial level of expected uncertainty–that is, how uncertainty interactions influence place cell mapping of environments.

To examine the consequences of uncertainties, we built a virtual reality spatial navigation task to test explicitly the impact of different levels, types, and interactions of variability on the cognitive map. As reward has been observed to be a particularly significant aspect of experience, potentially acting as an anchor for cognitive maps (Burgess and O’Keefe, 1996; Dupret et al, 2010; Gauthier and Tank, 2018; Jarzebowski et al, 2022; Sarel et al, 2017), we designed our task around uncertainty in the location of reward. Mice were trained, in a stable spatial reference frame, to lick for a water reward whose precise location on any trial was more or less certain (a form of expected uncertainty), and whose location distribution might also translate without warning (unexpected uncertainty). During the task, we imaged dorsal CA1 in the hippocampus using 2-photon calcium imaging (2P) (Dombeck et al, 2010) of pyramidal cells expressing the calcium indicator jGCaMP8m (Zhang et al, 2023). We found that expected uncertainty in reward location enhanced the proportion of place cells that tracked the reward on a trial-by-trial basis compared to what we refer to as low uncertainty. Additionally, the reward and the track end became anchors of a warped metric for space. Unexpected uncertainty caused substantial remapping of place cells but, when we varied the initial level of expected uncertainty, we did not find a difference in the overall proportion of place cells that remapped in the spatial reference frame. Instead, starting from a state of high versus low expected uncertainty increased the proportion of reward and warped place cells that moved to follow the reward after the unexpected change in reward location, a condition that we termed uncertainty interaction. Starting from a state of low expected uncertainty, by contrast, led to a less flexible representation in which reward location encoding place cells tended to remain at the location of the initial reward, even after the unexpected change in reward location. Hence, by inducing different forms of uncertainty in reward location and looking at their interaction, we show that uncertainty generates remapping in parallel, task-relevant, reference frames.

## Results

### Mice adapt their behaviour to the degree of uncertainty of the task

To study how different forms of reward location uncertainty affect the place cell code, we trained seven male, water-scheduled mice to lick for a water reward as they ran on a 3m linear virtual reality (VR) track, and simultaneously recorded place cell activity with 2-photon calcium imaging (Dombeck et al, 2010). At the end of the track, the screen switched off for 3 seconds and mice were teleported to the start of the track for the next trial. On each trial, the reward location lay at discrete sites within a designated reward zone of the track, with the width of this zone inducing varying degrees of predictability. In one subgroup (3 mice, low uncertainty; LU), the reward was made available at one of two adjacent locations 10 cm apart, generating low (but, to avoid potential anomalies, not zero) uncertainty about the reward location in each trial (Figure 1a:i). In the second group (4 mice, expected uncertainty; EU), the reward was made available at one of ten potential locations within a 1-meter zone, creating a condition in which mice could come to expect the resulting high aleatoric uncertainty (Figure 1a:ii). Importantly, the visual environments were the same between the two groups, and contained extra-track cues, as well as a more marked cue indicating the end of the track (Figure 1d). Once the mice were accustomed to the reward contingencies in low or expected uncertainty conditions, they all experienced a switch in reward location to a new, distal, 10 cm reward zone (Figure 1b). In the mice trained under LU, this switch induced a form of unexpected uncertainty (UU, Figure 1b:i). In the mice trained under EU, this switch induced a form of uncertainty interaction (UI, Figure 1b:ii).

**Fig. 1:**
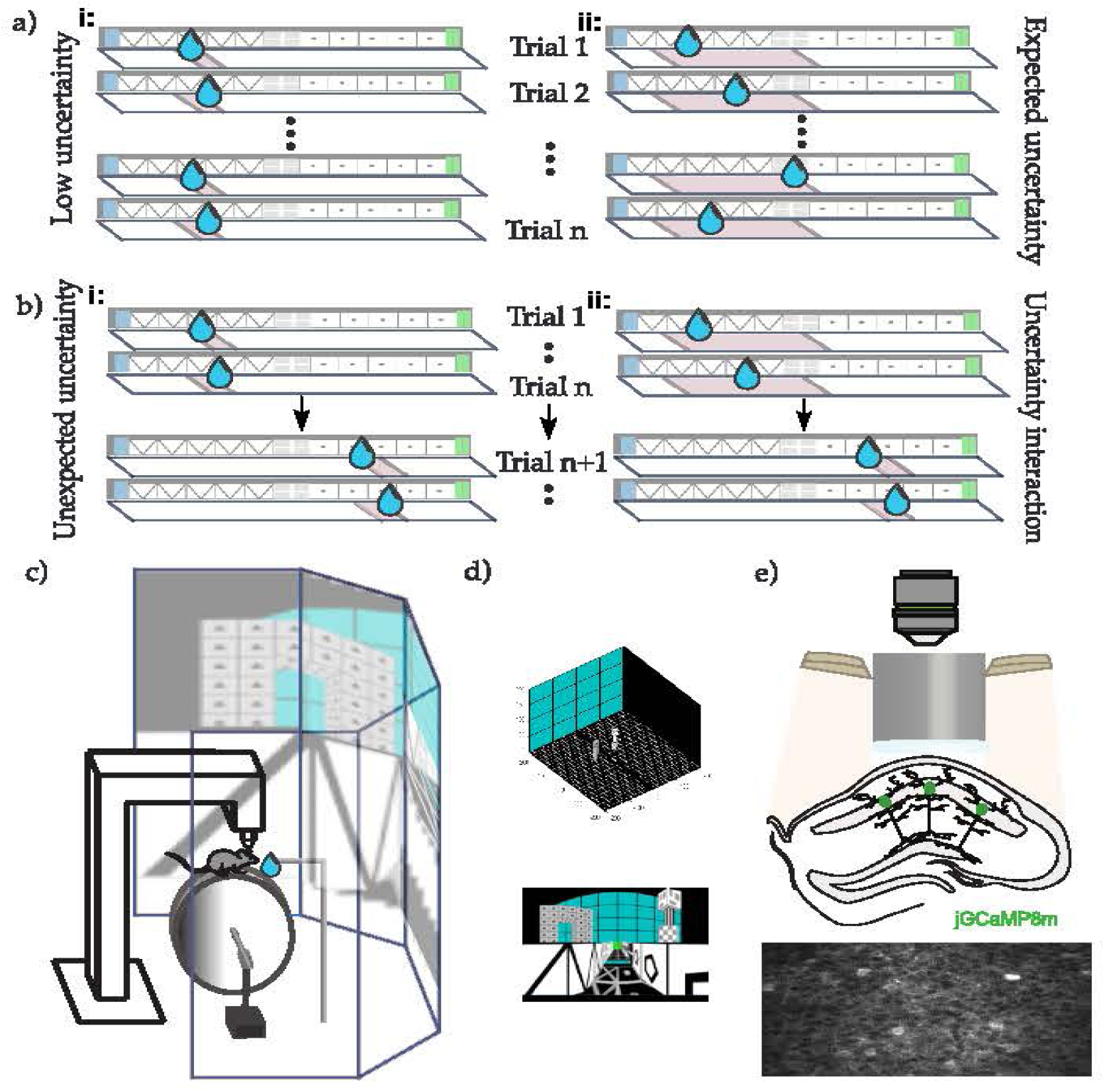
Training protocol and imaging procedure. a) Training protocol: head-fixed mice on a wheel ran in 1d virtual reality (VR) environments in which water reward was delivered at specific potential locations once per traversal of a 3m long linear track (and could subsequently be consumed anywhere by licking). In the low uncertainty condition (LU), the location could take one of two positions at the edges of a 10cm reward zone (left). In the expected uncertainty condition (EU), there were 10 potential locations evenly spaced within a 90cm wide zone that were selected uniformly at random on every run (right). Mice were trained on one session per day (on average 88.8 trials *±*15 std) until their behaviour was stable. b) After training, mice experienced a switch session. Initial trials (on average 40.8 *±*-3.5 std) in the session had the same location contingencies as those experienced during training. Without prior notice, the locations at which reward might be provided switched to one of two positions at the edges of a more distal 10cm zone, thus creating unexpected uncertainty (UU, left) in mice originally trained in LU, and a form of uncertainty interaction (UI, right) for mice originally trained in EU. c) Schematic of the VR apparatus: the licking behavior of mice was recorded as they ran on a wheel whose turning determined the velocity of the visual flow on screens. When the mice reached the end of the track, the screen went black for 3 seconds and mice were teleported to the start of the virtual track. d) Visualization of the track used for VR in this paper. Top: 3D view of the track, showing the relative perspective with distal cues. Bottom: front view of the track. e) Schematic of two photon calcium imaging of mouse CA1 neurons (green colors) expressing jGCaMP8m.

Mice were first trained until they were accustomed to the specific reward contingencies of LU and EU. Mice experiencing LU displayed licking and velocity patterns characteristic of high predictability: the lick rate increased shortly before the reward zone, peaked within it, and then decreased, stopping until the next trial (Figure 2a). These mice also slowed down as they approached the reward zone, stopped to consume the reward, and then resumed running at a faster velocity until the end of the track (Figure 2a).

**Fig. 2:**
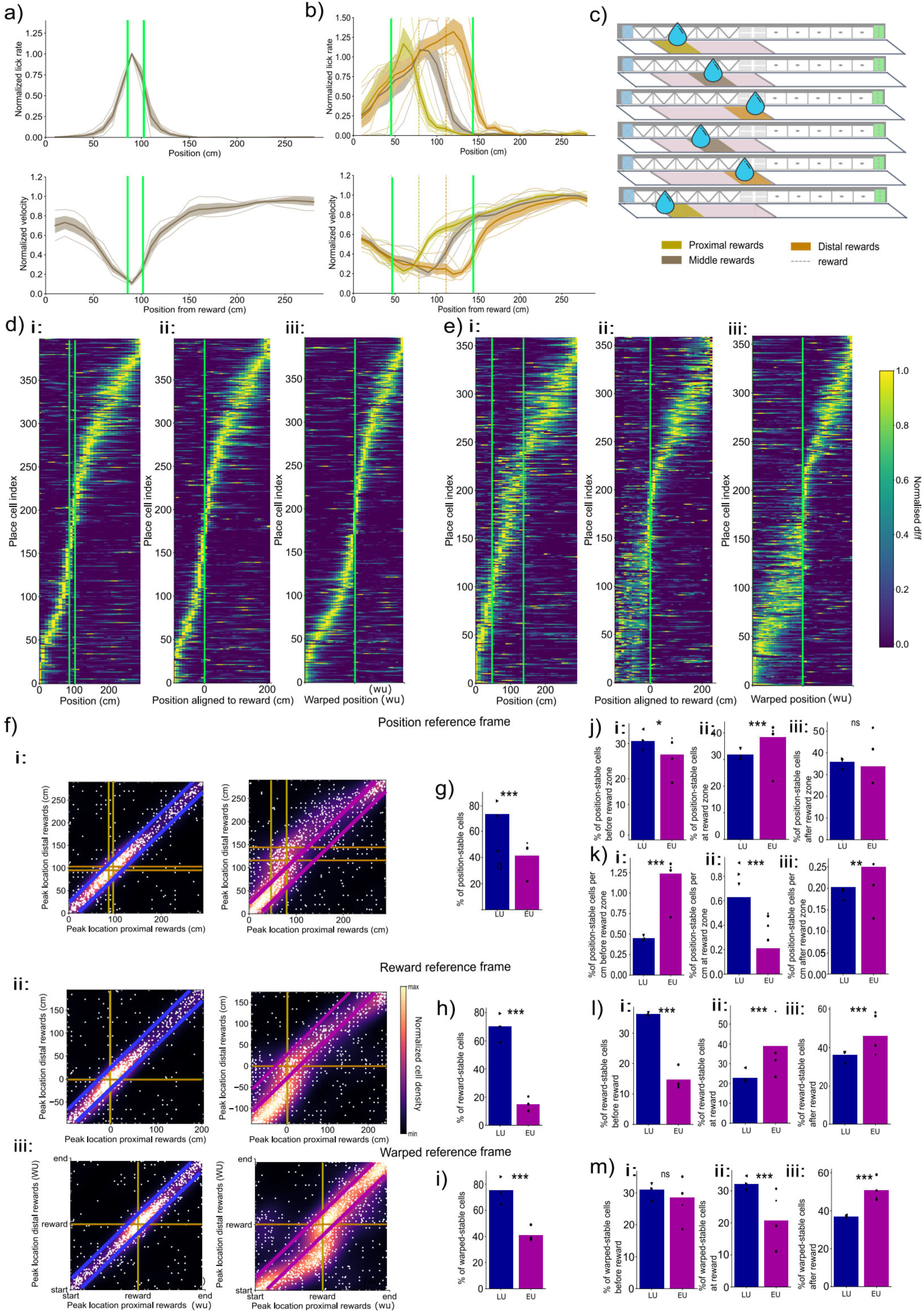
Expected uncertainty reveals dual spatial and reward reference frames for behaviour and place cell activity, and a warped metric that combines both. a) Top: Average lick rate (number of lick events per 10cm position bin) after training in the LU condition. Bottom: Average velocity trace in the same condition. For both: Thick line shows the mean across sessions (*n* = 12 sessions, *m* = 3 mice) normalised to the maximum per session, shaded region represents the standard deviation across sessions; shaded lines show each session trace. b) The same plots as for LU, but under EU, for laps of trials separated as shown in c): yellow: proximal, grey: middle and orange distal reward trials. For both top and bottom: Thick line showing the mean across sessions (*n* = 16 sessions, *m* = 4 mice) normalised to the maximum per session, shaded region represents the std across sessions, shaded lines show each session trace. Green thick lines show the reward zone. b) Diagram of the division between the laps according to the location at which the reward was consumed for the analysis in EU. c) i: Cross validated place map in a position reference frame for one session of LU for one animal, showing the average place cell activities (*N* = 437 place cells out of 518 total cells) on even trials normalised to their maximum value, ordered by their position of peak activity on odd trials, after training in low uncertainty. ii: The same activity, but averaged according to a reward reference frame (aligning the position to the reward location at every trial – see Methods). iii: The same activity averaged according to a warped/interpolated position-reward reference frame (a warped metric vector is created by two uniform interpolations linking the start of the track - reward - end of the track – see Methods). d) The same (d), but for an animal experiencing EU (*N* = 369 place cells out of 475 total cells). e) i: Scatter plot showing the positions of peak activity on trials on which the reward is at the proximal (x-axis) versus distal (y-axis) end of the reward zone for LU (left; 1118 place cells) and EU (right; 1192 place cells). Each white dot is a single place cell; the heatmaps show a probability density function estimate of the data (see Methods, normalised to 1). Yellow lines show the reward zone on proximal trials, orange lines on distal trials. Blue lines (left) and purple lines (right) delineate the diagonal used in the quantification for statistics in g). Scatter plots include a jitter proportional to cell density, enhancing visualization of overlapping data points. ii: Similar to (i), but in a reward-centered reference frame. The yellow line shows the reward location on proximal trials, the orange line on distal trials (both at 0, by definition of the reward reference frame). Blue square (left) and purple square (right) delineate the area used in the quantification for statistics in h). iii: similar to (i) in a warped metric (see Methods). Blue lines (left) and purple lines (right) delineate the post-reward diagonal used in the quantification for statistics in i). f) Percentages of cells that have a similar (*±*15cm) position of peak activity in ‘proximal’ and ‘distal’ reward trials in LU (blue region) and EU (purple region). comparison proportion z-test LU/EU *p* = 1 *×* 10*^−^*^53^, 1-sided proportion z-test LU*>*EU *p* = 5.26 *×* 10*^−^*^54^. g) Same than g) in a reward reference frame. Comparison proportion z-test LU/EU *p* = 1.1 *×* 10*^−^*^159^, 1-sided proportion z-test EU*>*LU *p* = 5.51 *×* 10*^−^*^160^. h) Same than g) in a warped reference frame (*±*3warped units). Comparison proportion z-test LU/EU *p* = 1.2 *×* 10*^−^*^61^, 1-sided proportion z-test EU*>*LU *p* = 5.6 *×* 10*^−^*^62^. i) Percentages of cells that are stable in a position reference frame (with a maximum displacement of *±*15cm; within the diagonal lines in e:i): i: before the reward zone, comparison proportion z-test *p* = 1.77 *×* 10*^−^*^2^, 1-sided proportion z-test LU*<*EU *p* = 8.86 *×* 10*^−^*^3^; ii: in the vicinity of the reward zone (−15 +20cm), comparison proportion z-test *p* = 8.82 *×* 10*^−^*^28^, 1-sided proportion z-test LU*>*EU *p* = 4.41 *×* 10*^−^*^28^, iii: after the reward zone, comparison proportion z-test *p* = 3.17 *×* 10*^−^*^2^, 1-sided proportion z-test LU*<*EU *p* = 1.59 *×* 10*^−^*^2^, for LU (blue) and EU (purple). j) Same as (j) but divided by the total area covered by every zone. left: comparison proportion z-test *p* = 3.47 *×* 10*^−^*^28^, 1-sided proportion z-test LU*<*EU *p* = 1.73 *×* 10*^−^*^28^ middle: comparison proportion z-test *p* = 1.96 *×* 10*^−^*^10^, 1-sided LU*>*EU comparison test *p* = 5.31 *×* 10*^−^*^11^, right: comparison proportion z-test *p* = 2 *×* 10*^−^*^1^. k) Percentages of cells with stable peaks in a reward reference frame (*±*15cm; within the boxes in e:ii): i: before the reward, comparison proportion z-test LU/EU *p* = 3.06 *×* 10*^−^*^8^, 1-sided proportion LU*>*EU z-test *p* = 1.53 *×* 10*^−^*^8^. ii: in the vicinity of the reward zone ([*−*15, +20]cm), comparison LU/EU proportion z-test *p* = 1 *×* 10*^−^*^5^, 1-sided proportion z-test LU*<*EU *p* = 5.02 *×* 10*^−^*^6^. iii: after the reward zone, comparison proportion z-test *p* = 1.27 *×* 10*^−^*^2^, 1-sided proportion z-test LU*<*EU *p* = 6.34 *×* 10*^−^*^3^, for LU (blue) and EU (purple). l) Percentages of cells with stable peaks in a warped reference frame (with a maximum displacement of 3 warped units, representing between 20cm and 40cm, depending on the position of the reward; within the diagonal lines in e:iii): i: before the reward, comparison proportion z-test LU/EU *p* = 3.37 *×* 10*^−^*^1^, non significant. ii:: in the vicinity of the reward zone (−2 +3 warped units), comparison proportion z-test *p* = 9.06 *×* 10*^−^*^6^, 1-sided proportion LU*>*EU z-test *p* = 4.53 *×* 10*^−^*^6^. iii: after the reward zone, for LU (blue) and EU (purple), comparison proportion z-test LU/EU *p* = 8.4 *×* 10*^−^*^7^, 1-sided proportion LU*<*EU z-test *p* = 4.2 *×* 10*^−^*^7^.

Mice trained in EU began licking and slowing down shortly before the start of the wider reward zone (Figure 2b) and therefore appeared to treat the reward as occurring anywhere across the broad zone, as expected for mice experiencing EU. To assess if the behavior in EU varied when the reward was consumed at different locations in the reward zone, we averaged the licking and velocity profiles over trials according to where the reward was consumed (see Methods) in the first third (proximal reward trials; 146 in total), middle third (115 middle reward trials), and last third (111 distal reward trials) of the reward zone. We found that mice licked persistently until they received the reward (Figure 2b). Once they received the reward, mice ceased licking and began running in a stereotypical manner (similar to LU; Figure 2b), demonstrating their understanding of the single-reward trial structure.

### Hippocampal place cells organise into position, reward-centred, and warped reference frames to reflect uncertainty

In order to investigate the population of place cells under these various conditions, we performed 2-photon calcium imaging of dorsal CA1 pyramidal cells while mice performed the task. We extracted place cells using an information theoretic criterion (see Methods), resulting in 1108 place cells for LU and 1192 place cells for EU. We first confirmed that the LU condition of our task produced results that were consistent with existing literature on place cell activity in reward navigation tasks by averaging place cell activity (DF/F) over the recording session post-training in an external, position, reference frame (Figure 2d:i). In the LU condition, we observed a higher density of place cells in the vicinity of the reward zone (Figure **??**:i), with on average 0.65% of cells per cm peaking in the vicinity of the reward zone defined as being between 15 cm before, and 20 cm after, it (see Methods) -, against 0.26% elsewhere (comparison in/out proportion z-test: p-value= 1.5 *×* 10*^−^*^75^, comparison in*>*out 1-sided proportion z-test: p-value*<* 7.3 *×* 10*^−^*^76^). Place cells peaking in the vicinity of the reward zone had narrower place fields (Figure **??**:iii; comparison in/out t-test: p-value= 1.8 *×* 10*^−^*^28^, comparison in*<*out 1-sided t-test: p-value= 8.9 *×* 10*^−^*^29^).

In the EU condition, we found only minor over-representation of the broad reward zone, with 0.35% of place cells per cm peaking in the region between *−*15cm of the start, and +20cm of the end, of the zone (see Methods), against 0.32% elsewhere (Figure **??**:i, comparison in/out proportion z-test: *p* = 1.3*×*10*^−^*^1^, EU comparison in*>*out 1-sided proportion z-test: *p* = 6.6 *×* 10*^−^*^2^). Place cells peaking in the vicinity of the reward zone were also narrower in EU (Figure **??**iii ; comparison in/out t-test: *p* = 1 *×* 10*^−^*^7^, comparison in*>*out 1-sided t-test: *p* = 5.18 *×* 10*^−^*^8^).

In this position reference frame, a higher proportion of cells was found to peak in the vicinity of the reward zone in LU than EU (Figure **??**i; comparison LU/EU proportion z-test: *p* = 7.9 *×* 10*^−^*^27^, comparison LU*>*EU 1-sided proportion z-test: *p* = 4 *×* 10*^−^*^27^).

To explore further the impact of a dynamically changing reward location on the place cell population on a trial-by-trial basis, we compared the positions of peak activity for each cell between trials in which the reward was collected near the start (proximal) or the end (distal) of the reward zone (scatter plots in Figure 2f:i; quantification of stable neurons in Figure 2g, whose peaks are within the bounds shown in Figure 2f:i). In the LU condition (Figure 2f:i; g:blue bar), in which these positions are very close, 73.10% of place cells maintained their peak activity location across the two groups of trials, compared to only 41.19% under EU (Figure 2f:i; g:purple bar; proportion z-test between the percentages in LU in EU: *p* = 1 *×* 10*^−^*^53^. 1-sided proportion z-test LU*>*EU *p* = 5.26 *×* 10*^−^*^54^). We also examined the locations of the peaks of the place fields of these cells and found that position-stable cells are evenly distributed, with 38.3% of those cells located before the reward zone for LU (c.f. 31.9% for EU), 27.9% (c.f. 32.22% for EU) in the vicinity of the reward zone, and 33.8% (c.f. 35.93% for EU) after the reward zone (Figure 2j). Although the total percentages are similar, the reward zone area is wider, and starts earlier in the track, in EU, resulting in a higher relative proportion of cells per cm before the reward zone (Figure 2k:i; comparison proportion z-test LU/EU *p* = 3.47 *×* 10*^−^*^28^, 1-sided proportion z-test LU*<*EU *p* = 4.41 *×* 10*^−^*^28^) and lower in the vicinity of the reward zone compared to LU (Figure 2k:ii; comparison proportion z-test LU/EU *p* = 8.82 *×* 10*^−^*^28^, 1-sided comparison z-test LU*>*EU *p* = 4.41 *×* 10*^−^*^28^).

Given that the reward changes location on a trial-by-trial basis, particularly in the EU condition, and that place cells can become organised within different task-relevant reference frames with experience (Anderson and Jeffery, 2003; Aoki et al, 2019; Gauthier and Tank, 2018; Markus et al, 1995; Muzzio et al, 2009; Plitt and Giocomo, 2021; Radvansky et al, 2021; Sosa and Giocomo, 2021; Sosa et al, 2023), specifically reward (Burgess and O’Keefe, 1996; Gauthier and Tank, 2018; Jarzebowski et al, 2022; Sosa and Giocomo, 2021; Sosa et al, 2023), we asked whether the EU condition might reinforce the reward reference frame, possibly reflected in an increased population representing the changing variable. We therefore considered whether cells code for position *relative* to the reward location on a trial rather than in spatial position associated with the track. To examine this we averaged cell activity relative to reward position (Figures 2d:ii; e:ii, see Methods). In contrast to the position reference frame, in the reward reference frame, there was an equal accumulation of cells aligned in the vicinity of the reward in both LU and EU conditions (Figure **??**ii), with 4.8% of cells per cm in the vicinity of the reward (with a peak of activity between *−*15cm and +20cm of the reward), against only 0.2% of cells per cm outside these bounds, in LU (comparison in/out proportion z-test: p-value*<* 0.2 *×* 10*^−^*^308^, comparison in*>*out 1-sided proportion z-test: p-value*<* 2.2 *×* 10*^−^*^308^), and 3.4% per cm in the vicinity of the reward, against 1.25% per cm elsewhere in EU (comparison in/out proportion z-test: p-value*<* 2.2 *×* 10*^−^*^308^, comparison in*>*out 1-sided proportion z-test: p-value*<* 2.2 *×* 10*^−^*^308^; Figure **??**ii). Averaging in a reward-centered reference frame also reduced the widths of place fields peaking in the vicinity of the reward compared to elsewhere, for both LU (comparison in/out proportion z-test: p-value= 3.2 *×* 10*^−^*^28^, comparison in*<*out 1-sided proportion z-test: p-value*<* 1.61 *×* 10*^−^*^28^) and EU (comparison in/out proportion z-test: p-value= 1.6*×*10*^−^*^12^, comparison in*<*out 1-sided proportion z-test: p-value*<* 8.11*×*10*^−^*^13^; Figure **??**iv). The difference in accumulation of reward place cells between position and reward reference frames revealed a population of cells that stably followed the reward on every trial (termed reward cells) and was confirmed by single-cell activity profiles across all trials (Figure **??**), highlighting populations of cells with stable fields relative to position and also reward. These reward cells generalize previous findings (Gauthier and Tank, 2018) to our task in which the reward changes location on every trial.

To investigate the effect of a dynamically changing reward location on the place map in this reward reference frame, we compared the locations of peak activity with respect to reward location between proximal and distal reward trials. We found that 70.21% of the place cells maintained their peak activity relative to the reward location across the two groups of trials in LU, compared with 14.85% in EU (Figure 2f:ii, with the stable neurons shown in the boxes quantified in Figure 2h; proportion z-test LU/EU: *p* = 1 *×* 10*^−^*^159^, 1-sided proportion z-test LU*>*EU *p* = 5.51 *×* 10*^−^*^160^). Examining the distribution of these cells along the track (Figure 2l), we found that 36.25% of the reward-stable cells were before the reward in LU, compared to 14.69% in EU (Figure 2l:i; comparison proportion z-test *p* = 3.06 *×* 10*^−^*^8^, 1-sided proportion LU*>*EU z-test *p* = 1.53 *×* 10*^−^*^8^). Stability of encoding at the reward was most enhanced in EU, with 38.98% of reward-stable cells being located in its vicinity, compared to 22.87% of reward-stable cells in LU (Figure 2l:ii; comparison LU/EU proportion z-test *p* = 5.02 *×* 10*^−^*^5^, 1-sided proportion z-test LU*<*EU *p* = 5.02 *×* 10*^−^*^6^). After the reward, less stability was reported in LU, with 36.24% of reward-stable cells compared to EU, with 46.33% of reward-stable cells (Figure 2l:iii; comparison proportion z-test *p* = 1.27 *×* 10*^−^*^2^, 1-sided proportion z-test LU*<*EU *p* = 6.34 *×* 10*^−^*^3^).

While the reward location cannot be predicted, the part of each run from the reward to the end of the track is predictable, and is characterized by a stereotypical behavioral routine. Hippocampal activities have been shown to reflect the statistics of the episodic environments animals experience (Plitt and Giocomo, 2021), for example reflecting stereotypical behavioural sequences (Skaggs and McNaughton, 1998), and organising along warped metrics that homogenise similar episodes (Gothard et al, 1996). We therefore asked whether the hippocampus might similarly represent these post-reward events regardless of reward location, reflecting stereotypical changes. Consistent with this idea, we qualitatively observed a group of cells that seemed to span the range from the reward location to the end of the track in a flexible manner (Figure 2f:i). To quantify this, we considered a third, warped, metric for space in which we compressed and expanded it so that there were two segments of 20 bins each — one linking the start of the track to the reward location and the other from the reward location to the end of the track (Figure 2d:iii; e:iii). We found that 75.4% of cells kept their position of peak activity in the warped reference frame between proximal and distal reward trials in LU, and 41.1% in EU (Figure 2i). Examining the relative distributions of warped-stable cells around the reward location in this reference frame, we found a balanced distribution in LU: 31.01% of warped-stable cells before the reward, 32.0% in the vicinity of the reward, and 36.8% after the reward (Figure 2m). As the reward location cannot be predicted in EU, this analysis is provided here for completeness with respect to other reference frames. In contrast, the warped metric highlighted a post-reward alignment of cells after the reward in EU, with 28.51% of the warped-stable cells before the reward, 20.7% in the vicinity of the reward, and 50.7% after the reward. We found a significantly lower degree of post-reward warping in LU compared to EU (Figure 2m:iii; comparison proportion z-test *p* = 8.40 *×* 10*^−^*^7^, 1-sided proportion z-test LU*<*EU *p* = 4.2 *×* 10*^−^*^7^).

Note that the higher percentages of stability between proximal and distal reward trials reported in Figures 2g,h,i in low uncertainty simply reflect the task design, in which reward, position and warped reference frames are more similar in LU than in EU due to its far narrower reward zone. We provide the statistical comparisons for the sake of completeness. Our finding of excess stability in reward (Figure 2l) and warped reference frames (Figure 2m) in EU confirm our conclusion that expected uncertainty highlights enhanced reward and warped reference frames as an adaptation to reflect the statistics of change in the task design. Overall, our findings show that expected uncertainty in reward location enhanced the proportion of place cells that tracked the reward on a trial-by-trial basis (reward-referenced cells) compared to low uncertainty, and the reward and the track end became anchors of a warped metric for space.

### Expected uncertainty leads to enhanced flexibility of the reward and warped populations towards a surprisingly new reward location

We have so far shown that expected uncertainty leads to an enhanced reward and warped reference frame by contrasting with a condition of low uncertainty. To complement the collection of uncertainties, and investigate their interaction, we performed a larger, unpredictable, change in reward location meant to induce a sudden surprise and a condition of unexpected uncertainty. After familiarising the animals to the LU and EU conditions, we imaged CA1 place cells while changing the reward location during a session, without prior notice or experience, to a narrow reward zone further down the track (Figure 1b). This unannounced change generates unexpected uncertainty (UU) in LU mice and a form of uncertainty interaction (UI) in EU mice. As drastic changes in context can lead to very abrupt shifts in place field locations (Michon et al, 2021; Sheffield et al, 2017; Wills et al, 2005), we asked whether UU would induce a higher degree of change in the place map compared to UI, due to a higher level of surprise. Comparison between LU and EU highlighted a difference with respect to the anchor of the reward on the place map, but it was unclear if or how these differences would change the response to unexpected uncertainty. Specifically, noting that the reward location variability in the EU mice led them to have a greater proportion of place cells stably tied to reward and warped reference frames rather than position, we tested whether this would generalize to the farther move of the reward, which would be exemplified by greater stability in these reference frames for UI than UU in the face of an unexpected change.

We first verified that the behavior after the switch stabilizes to a pattern reflecting the new task statistics. The licking and velocity patterns aligned with prior observations (compare Figures 2a;b and 3a;b). Note that the two subjects in UI had different patterns of post-switch behavior (Figure **??**); thus, as well as analyzing them together we report in the Supplement (Figures S**??** and S**??**) the statistical comparisons presented in this section for each of these animals separately.

Next, we examined how the place map was impacted by the unexpected change. Although we expected more global remapping in UU than UI (since they should have been more surprised), a qualitative assessment of the place map after the switch (Figure 3c;d) highlighted similar, moderate, degrees of remapping in both conditions, primarily affecting place cells peaking between the previous and new reward zones. Comparing post-switch maps, we found that reward over-representation was marginally less in the new location under UU than UI. Indeed, after UU we found 0.38% of cells per cm in the vicinity of the reward zone, 0.32% of cells per cm elsewhere, versus 0.56% of cells per cm in the vicinity of the reward after UI and 0.28% of cells per cm elsewhere (proportion z-test UU/UI *p* = 1.13 *×* 10*^−^*^2^, 1-sided z-test UU*<*UI *p* = 5.6 *×* 10*^−^*^3^, Figure **??**).

**Fig. 3:**
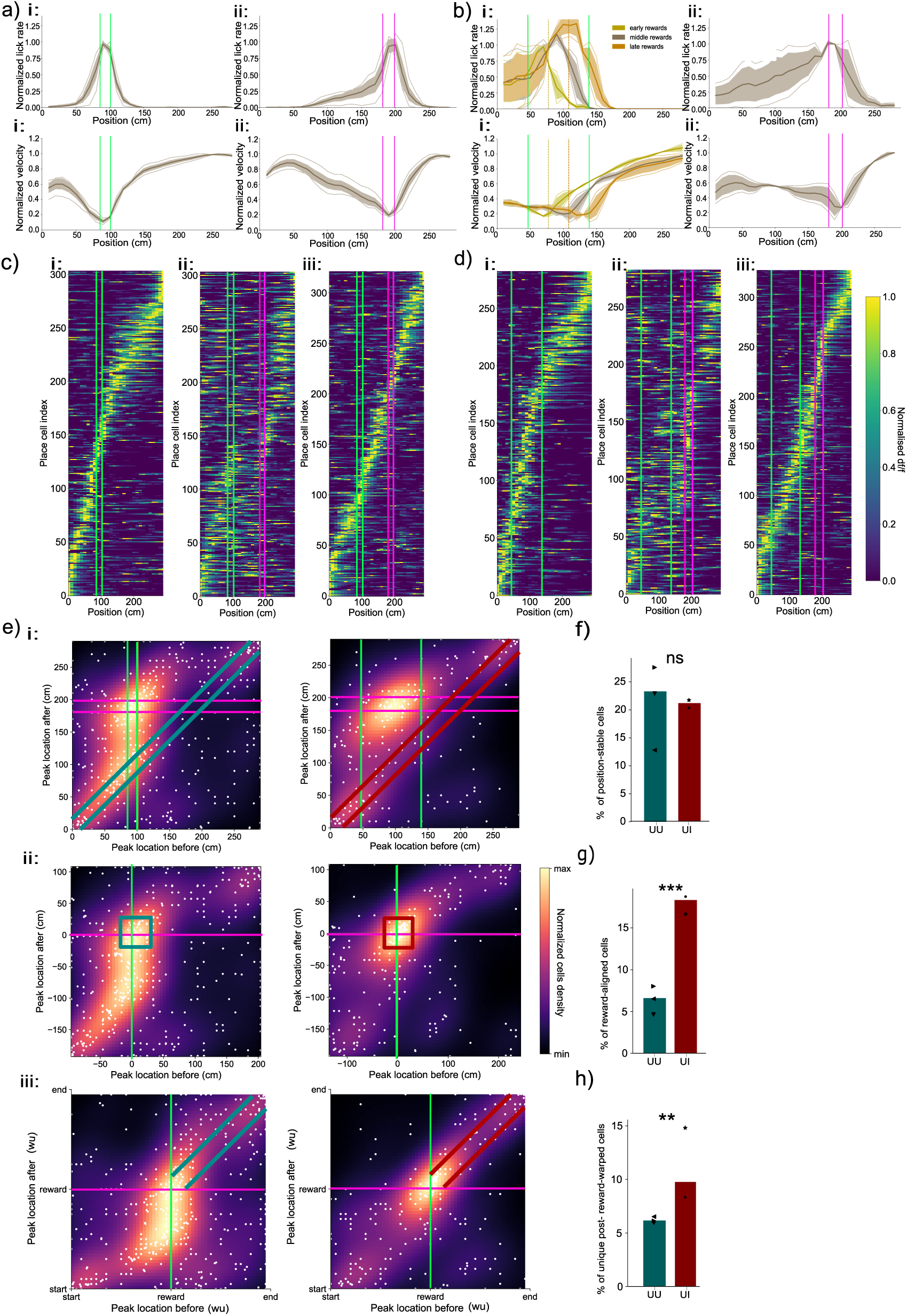
Expected uncertainty in reward location enhances flexible reward and warped reference frames. a) UU: Top: normalized lick rate averaged over all sessions. i: before the switch, ii: after the switch. Bottom: similar to Top) for normalized velocity. Green thick lines show the full reward zone. Shaded areas show standard deviations and individual lines show individual sessions averages. b) UI: Top: normalized lick rate on proximal (Yellow), middle (Grey) and distal (Orange) reward trials. i: before the switch, ii: after the switch. Bottom: similar to Top) but for normalized velocity. Green thick lines show the full reward zone before the switch, pink lines the reward zone after the switch. Shaded areas show standard deviations and individual lines show individual sessions averages. See Figure **??** for results for separate mice. c) i: Place map before the switch (N place cells=304) in UU, showing the average activity for one animal, ordered according to their cross-validated position of peak activity before the switch, and shown in a position reference frame. Green lines mark the reward zone. ii: activities of the same cells ordered as in (i), after the switch. Turquoise lines mark the previous reward zone, pink lines show the new reward zone. iii: New place map after the switch (N after=322). d) The same as (c), but for UI (N before=283, N after=328). e) i: Scatter plot showing the positions in a position reference frame of peak activity before (x-axis) versus after the switch (y-axis) in UU (left) and UI (right). Each white dot is a cell and heatmap shows a probability density function estimate (see Methods). Turquoise lines delineate the diagonal used for statistics in f). Scatter plots include a jitter proportional to cell density, enhancing visualization of overlapping data points. ii: Similar to i: but in a reward reference frame. Turquoise squares delineate the area used for statistics in g). iii: Similar to (i:,ii:) but in a warped reference frame. Turquoise lines delineate the post-reward diagonal used for statistics in g). f) Percentages of cells with stable peaks in a position reference frame (with a maximum displacement of *±*15cm; shown by the lines in the heatmap in e)i:) : for UU (turquoise) and UI (red). The black dots show individual session percentages. Comparison proportion z-test UU/UI *p* = 5.14 *×* 10*^−^*^1^. g) Percentages of cells with stable peaks in a reward reference frame (between *−*15cm and +20cm of the reward; shown by the lines in the heatmap in e)ii:), excluding position-stable cells. Proportion z-test UI/UU *p* = 1.73 *×* 10*^−^*^6^, 1-sided comparison UU*<*UI 1-sided proportion z-test *p* = 8.65 *×* 10*^−^*^7^. h) Percentages of cells with stable peaks in a warped reference frame (with a maximum displacement of 3 warped units, representing between 20cm and 40cm, depending on the position of the reward, shown by the lines on the heatmaps in e)iii:), excluding position- and reward-stable cells. Proportion z-test UU/UI *p* = 2.94 *×* 10*^−^*^3^, 1-sided comparison UU*<*UI 1-sided proportion z-test *p* = 1.47 *×* 10*^−^*^3^.

To quantify the impact of the sudden reward location change on each place cell’s activity, we compared the location of peak activity before and after the switch for cells that remained place cells after the switch (455 place cells out of 1872 total cells for UU, 246 out of 970 for UI). Surprisingly and contrarily to our expectations, we found that similar percentages of cells maintained their peak activity location after the switch in UU (24.4% of cells) and UI (21.5% of cells) in a position reference frame (Figure 3e;h; comparison proportion z-test UU/UI *p* = 4 *×* 10*^−^*^1^). Thus, unexpected uncertainty caused substantial remapping of place cells but, when we varied the initial level of expected uncertainty, we did not find a difference in the overall proportion of place cells that remapped in the spatial reference frame.

However, building on the observation of a slightly lower reward over-representation after UU compared to UI, we turned to analyse the cells that moved with the reward across the switch in a reward reference frame. We found that a significantly lower percentage of cells stably peaked in the vicinity of the reward in a reward-reference frame across the switch in UU (7% of cells) than in UI (18.2% of cells) (Figure 3f;i; comparison proportion z-test UU/UI *p* = 1.73 *×* 10*^−^*^6^, 1-sided proportion z-test UU*<*UI *p* = 8.65 *×* 10*^−^*^7^), excluding those place cells that were stable in the position reference frame (i.e., those quantified in Figure 3h). Thus, expected uncertainty leads to a more flexible reward reference frame.

We then wondered whether the enhanced flexibility of the reward anchor in UI would also translate to the warped reference frame. Indeed, we found that fewer cells maintained their peak activity in UU (6% of cells) compared to UI (14% of cells) in the warped reference frame (Figures 3g;j; comparison proportion z-test UU/UI *p* = 2.94 *×* 10*^−^*^3^, 1-sided proportion z-test UU*<*UI *p* = 1.47 *×* 10*^−^*^3^), excluding any position-stable or reward-peaking cells quantified in Figures 3h;i. Therefore, expected uncertainty leads to hippocampal representations that are more stable in both the reward and warped reference frames in subsequent adaptations to unexpected changes.

### Place cells over-represent previous rewards in UU, generalize in UI

Surpised by the finding that the proportion of cells maintaining their peak activity location before and after the switch was similar in UU and UI conditions in a position reference frame, we decided to investigate further the relative stability in position reference frame, and looked at how the peaks of these position-stable cells were distributed along the track. The distributions of percentage of cells per position bin did not show any significant overall difference between the two conditions (Kolmogorov-Smirnov test *p* = 0.872, non significant; figure 4a).

**Fig. 4:**
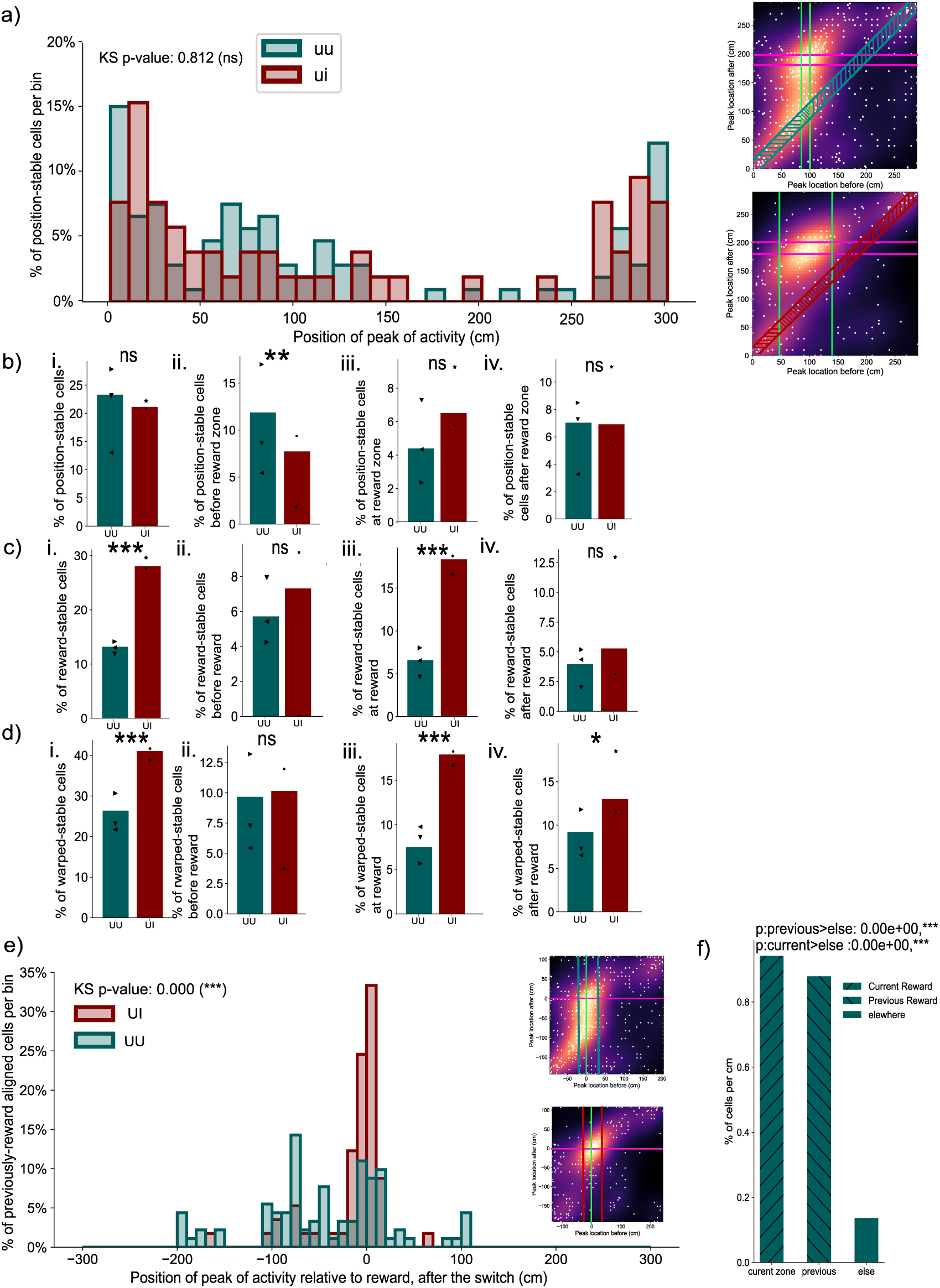
Unexpected uncertainty in reward location highlights persistence of the previous reward location, EU features generalisation of reward encoding. a) Distribution of the locations of the peak activity of position-stable cells. The x-axis shows position along the track in cm. Bars show the percentage of position-stable cells having their peak activity in a position reference frame in the respective position-bin for UI (red) and UU (turquoise). Right insets show repeat of 3e) illustrating the cells counted in the histogram plot. Distribution comparison using a Kolmogorov-Smirnov test *p* = 0.47, non significant. b) Percentages of cells with stable peaks in a position reference frame: i: across the whole track (with a maximum displacement of *±*15cm; similar to 3h) comparison UU/UI proportions z-test *p* = 5.14 *×* 10*^−^*^1^, non significant; ii: before the reward zone (*<−*15cm) before and after the switch (horizontal bar zone in the insert); comparison UU/UI proportions z-test *p* = 8.64 *×* 10*^−^*^2^, comparison UU*>*UI 1-sided proportion z-test *p* = 4.32 *×* 10*^−^*^2^; iii: in the vicinity of the reward zone (*−*15cm/+20cm) before and after the switch (tilted bar zone in the insert); comparison UU/UI proportions z-test *p* = 2.27 *×* 10*^−^*^1^, non significant; iv) after the reward zone (*>*+20cm) both before and after the switch (vertical bar zone in the insert); comparison UU/UI proportions z-test *p* = 9.52 *×* 10*^−^*^1^, non significant. c) Percentages of cells with stable peaks in a reward reference frame: i: across the whole track; comparison UU/UI proportions z-test *p* = 1.26 *×* 10*^−^*^6^, comparison UU*<*UI 1-sided proportion z-test *p* = 8.65 *×* 10*^−^*^7^; ii: before the reward zone (*<−*15cm) before and after the switch; comparison UU/UI proportions z-test *p* = 4.04 *×* 10*^−^*^1^, non significant; iii: in the vicinity of the reward (*−*15cm/+20cm) before and after the switch; comparison UU/UI proportions z-test *p* = 1.73 *×* 10*^−^*^6^, comparison UU*<*UI 1-sided proportion z-test *p* = 8.65 *×* 10*^−^*^7^; iv) after the reward (*>*+20cm) before and after the switch; comparison UU/UI proportions z-test *p* = 4.14 *×* 10*^−^*^1^, non significant. d) Percentages of cells with stable peaks in a warped reference frame: i: across the whole track; UU/UI proportions z-test *p* = 6.51 *×* 10*^−^*^5^, comparison UU*<*UI 1-sided proportion z-test *p* = 3.26 *×* 10*^−^*^5^; ii: before the reward (*<*-2 warped units) before and after the switch; comparison UU/UI proportions z-test *p* = 8.35 *×* 10*^−^*^1^, non-significant; iii: in the vicinity of the reward (*−*2*/* + 3 warped units) before and after the switch, comparison UU/UI proportions z-test *p* = 2.9 *×* 10*^−^*^5^, comparison UU*<*UI 1-sided proportion z-test *p* = 1.4 *×* 10*^−^*^5^; iv: after the reward (*>*+3 warped units) before and after the switch; comparison UU/UI proportions z-test *p* = 1.2 *×* 10*^−^*^1^, comparison UU*<*UI 1-sided proportion z-test *p* = 6.2 *×* 10*^−^*^2^. e) Distribution of the peak location relative to post-switch reward of the cells that peaked in the vicinity of ([*−*15, +20]cm) the reward before the switch for UI (red) and UU (turquoise). The x-axis shows bins of position along the track relative to post-switch reward. Cells peaking at 0cm follow the reward through the switch. Right insets repeat figure 3e), illustrating the cells counted in the histogram plot with vertical bars. Distribution comparison using a Kolmogorov-Smirnov test p-value*<* 2.2 *×* 10*^−^*^308^. f) Percentages per cm of previously reward peaking cells after the switch, that stay reward peaking (current, rightward tilt), that stay peaking at the previous reward (previous, leftward tilt), or that move elsewhere (else, plain bar) for UU. Comparison current*>*else 1-sided propor-tions z-test *<* 2.2 *×* 10*^−^*^308^, comparison previous*>*else 1-sided proportions z-test *<* 2.2 *×* 10*^−^*^308^.

However, minor differences were apparent when dividing the track up into three areas: ahead of the reward zone before the switch, at the previous reward zone, and the remainder (zones marked with bars in the insert in Figure 4a). In UU, a slightly greater position-stability was observed before the previous reward zone, with 11.87% of cells (50.5% of total position-stable cells) located before the previous reward zone in UU, compared to 7.72% of cells (35.8% of total position-stable cells) in UI (Figure 4b:ii; comparison UU/UI proportions z-test *p* = 6*×*10*^−^*^2^, 1-sided proportion z-test UU*>*UI *p* = 2.27*×*10*^−^*^1^). In the vicinity of the previous reward zone, no significant difference was found, with 4.4% of cells (18% of position-stable cells) in UU and 6.5% of cells (30% of position-stable cells) in UI (Figure 4b:iii; comparison UU/UI proportions z-test *p* = 2.2 *×* 10*^−^*^1^, non significant). After the previous reward zone, a similar proportion of cells was position-stable, with 7.7% of cells (31.5% of position-stable cells) in UU and 7.3% of cells (34% of position-stable cells) in UI (Figure 4b:iv; comparison UU/UI proportions z-test *p* = 8.6 *×* 10*^−^*^1^, non significant).

Consistent with our results so far, this picture changed considerably in a reward reference frame (Figure 4c). Here, we found greater overall reward-stability in UI, with only 13.6% of cells maintaining their peak activity location relative to the reward after the switch in UU, compared to 28% of cells in UI (Figure 4c:i; comparison UU/UI proportions z-test *p* = 3 *×* 10*^−^*^6^, comparison UU*<*UI 1-sided proportion z-test *p* = 2.7 *×* 10*^−^*^6^). Specifically, 5.4% of cells (40.3% of reward-stable cells) were located before the reward in UU, versus 7.3% of cells (26% of reward-stable cells) in UI (Figure 4c:ii; comparison UU/UI proportions z-test *p* = 3.4 *×* 10*^−^*^1^, non significant). In the vicinity of the reward, we found 7% of cells (51.1% of reward-stable cells) in UU, and 18.3% of cells (65.2% of reward-stable cells) in UI (Figure 4c:iii; comparison UU/UI proportions z-test *p* = 3.4 *×* 10*^−^*^6^, comparison UU*<*UI 1-sided proportion z-test *p* = 2.7 *×* 10*^−^*^6^). Post-reward, similar percentages of reward-stable cells were found, with 4.8% of cells (35.5% of reward-stable cells) in UU, and 5.3% of cells (18.8% of reward-stable cells) in UI (Figure 4c:iv; comparison UU/UI proportions z-test *p* = 7.9 *×* 10*^−^*^1^, non significant).

Given the over-representation of reward locations in both LU and EU, and our discovery that expected uncertainty leads to an enhanced flexibility of this population, we sought to understand where the previously reward-peaking cells moved to after the unexpected switch in UU and UI. For this, we explored the post-switch peak locations of cells that peaked in the vicinity of the reward pre-switch. These differed significantly between UU and UI (Figure 4e; distribution comparison using a Kolmogorov-Smirnov test p-value*<* 2.2*×*10*^−^*^308^). The bimodal distribution for UU indicates a peak at the previous reward location; by contrast, more reward-cells moved to the new reward location in UI. Quantifying whether the previous reward location in UU is indeed over-represented, we found 0.9% of previously reward-peaking cells per cm at the current reward location after the switch (comparison current*>*elsewhere 1-sided proportion z-test p*<* 2.2*×*10*^−^*^308^) and 0.88% of those cells per cm peaking at the previous reward location (comparison previous¿elsewhere 1-sided proportion z-test *<* 2.2 *×* 10*^−^*^308^), while only 0.14% of previously reward stable cells remapped elsewhere in the track after the switch (See figure 4f), confirming persistence of the previous reward location in UU.

Building on the observation of an enhanced post-reward warped metric after UI compared to UU, we turned to look into whether there was a difference between UU and UI in how the warped stability is organized with respect to the reward. Investigation of the stability with respect to the warped reference frame confirmed our earlier results, showing overall lower warped-stability in UU, with 26.2% of cells, compared to UI, with 41% of cells (Figure 4d:i; comparison UU/UI proportions z-test *p* = 6.51 *×* 10*^−^*^5^, comparison UU*<*UI 1-sided proportion z-test *p* = 3.26 *×* 10*^−^*^5^). Similarly consistent with our earlier results, no difference in warped-stability was found before the reward, with 10.1% (38.7% of all warped-stable cells) in UU and 10.2% (24.8% of warped-stable) in UI (Figure 4d:ii; comparison UU/UI proportions z-test *p* = 9.8 *×* 10*^−^*^1^, non-significant). In the vicinity of the reward, fewer cells were warped-aligned in UU (7.4% of cells, 28.6% of warped-stable cells), compared to UI (17.9% of cells, 43.6% of warped-stable cells) (Figure 4d:iii; comparison UU/UI proportions z-test *p* = 2.9 *×* 10*^−^*^5^, comparison UU*<*UI 1-sided proportion z-test *p* = 1.4 *×* 10*^−^*^5^). Post-reward warped-stability was slightly different, with only 8.6% of cells (32.8% of warped-stable cells) in UU, and 13% (31.7% of warped-stable cells) in UI (Figure 4d:iv; comparison UU*<*UI 1-sided proportion z-test *p* = 6.02 *×* 10*^−^*^2^).

Overall, these results establish that starting from a state of high versus low expected uncertainty increased the proportion of reward and warped place cells that moved to follow the reward after the unexpected change in reward location. Starting from a state of low uncertainty, by contrast, led to a less flexible representation in which reward location encoding place cells tended to remain at the location of the initial reward, even after the unexpected change in reward location.

## Discussion

We imaged dorsal CA1 while mice navigated in a virtual reality corridor in which reward became available according to one of a number of distributions of spatial location. These induced different forms of uncertainty that we studied across three positional reference frames: environment-centered, reward-centered, and a combined metric where the reward and the end of the track anchored experience, with the hippocampus generating what amounts to a warped spatial metric. We found that reward-dedicated place cells adapted flexibly to trial-by-trial changes in reward location, with this adaptability extending to larger, unexpected reward shifts, especially in reward-based and warped reference frames. This was not observed in animals conditioned to low uncertainty. Initial stability in reward location did not lead to more global remapping in a position reference frame when the reward subsequently moved, but led to persistence of previous reward location. These results contribute to our understanding of the structure of cognitive maps.

Our results expand on previous findings about reward-dedicated place cells(Dupret et al, 2010; Gauthier and Tank, 2018; Hollup et al, 2001; Jarzebowski et al, 2022; Sosa and Giocomo, 2021), showing their ability to adapt to single-trial changes in reward location. This is consistent with previous electrophysiological results highlighting abstract goal populations in the hippocampus (McKenzie et al, 2013; McNaughton and Bannerman, 2024; Zeithamova et al, 2018), and behavioral results showing that the hippocampus is required for single-shot learning of new goal locations (Bast et al, 2009; Kleinknecht et al, 2012; Morris et al, 1990; Sosa and Giocomo, 2021; Steele and Morris, 1999; Tessereau et al, 2021). Such cells have been suggested in models (Burgess and O’Keefe, 1996; Foster et al, 2000; Tessereau et al, 2021) as serving flexible behavioural adaptation, for example acting as a reference point for vector-based navigation (Burgess et al, 1995; Foster et al, 2000; Tessereau et al, 2021), or uncertainty resolution, to guide prediction (Burgess et al, 1995). Our results converge with a recent paper investigating the effect of multiple similar changes in reward location on the reward population codes of the hippocampus. By changing the reward location between multiple phase of stable reward locations, (Sosa et al, 2023) found that place cells can organise within reward-centered sequences which recruit more cells as the reward location changes day-by-day. Although the authors focus on reward population codes, we can now interpret their results in terms of expected uncertainty, induced by block-by-block changes in reward. The extra recruitment of reward cells would then be an instance of the excess of reward-following cells apparent in our UI condition. Similar findings suggest that reward-induced behavioral changes create a landmark-based reference frame in the hippocampus (Vaidya et al, 2023), with over-representation of salient cues extending beyond rewards (Tanni et al, 2022; Vaidya et al, 2023). This over-representation likely arises from distinct mechanisms for landmarks and rewards (Sato et al, 2020).

In conditions of EU, we observed a warped spatial metric consistent with past studies(Gothard et al, 1996), where the track segment following the reward and preceding the teleportation zone was renormalized. Whether the warped metric is the reflection of stereotypical behavioural sequences induced by having to stop to consume the reward, and running until the end of the track, or whether the reward itself is a sufficient anchor to induce such a warped metric, remains unclear. Comparable place map warping has been seen when mice were exposed to gradually changing visual patterns Plitt and Gio-como (2021) or visual boundaries (Leutgeb et al, 2005a), creating continuous place cell activity profiles. In contrast, abrupt remapping occurred when mice were only familiar with extreme conditions, paralleling the response to unexpected uncertainty in the reward reference frame in our study. The integration of homogeneous episodes within continuous, possibly warped, metrics is also consistent with suggested roles of the hippocampus as a comparator (Kumaran and Maguire, 2007; Vinogradova, 2001) – perhaps responding to the conflict between external cues and internal, self-motion cues (Gothard et al, 1996), or intrinsic reward encoding. Indeed, warped metrics provide an efficient way to associate discontiguous events (Wallenstein et al, 1998), and may promote one-shot decision making by enhancing state-space separability (McKenzie et al, 2014; Muzzio et al, 2009; Nitz, 2009; Sun et al, 2023).

Our finding that unexpected uncertainty did not induce greater position remapping than expected uncertainty contradicts our initial hypothesis, which anticipated more extensive remapping under surprise. By contrast, previous work has suggested that greater surprise is associated with greater remapping (Sanders et al, 2020), and indeed drastic changes in context, such as the visual environment (Anderson and Jeffery, 2003; Bostock et al, 1991; Kentros et al, 1998; Leutgeb et al, 2005b; Muller and Kubie, 1987; Sanders et al, 2020; Sheffield and Dombeck, 2019) can lead to substantial degrees of remapping. It may be that surprising reward locations and sensory mispredictions (Sanders et al, 2020) are treated somewhat independently. This would be consistent with the greater degree of reward-related and warped-metric remapping in UU compared to UI, suggesting that remapping can occur independently in different reference frames, and building on existing results shedding light on overlapping reference frames in spatial navigation tasks (Zinyuk et al, 2000).

In UU, we found that the population of place cells previously peaking at the reward became bimodally distributed around the previous and new reward location. This suggests that repeated experience of a specific episode could lead to cells becoming specific to single episodes, akin to splitter cells (Wood et al, 2000), but in reward reference-frames, similar to the finding in (McKenzie et al, 2013). In contrast, in UI, reward-aligned cells and warped-aligned cells moved flexibly to the new goal location. This confirms a previous result suggesting independence of reward and position reference frames in rats (Aoki et al, 2019). We might interpret this difference in terms of generalization: context-specific representations are probably well suited for efficient decision making when environments distinctly differ, as in the transition in UU. However, under EU, the multiple reward locations are tied under a common, moderately compact, distribution. Rather than exhausting capacity by representing each separately, the hippocampal solution appears to be to have similar events share representations, by adopting metrics that encapsulate shared aspects of experience. This then generalizes when the reward location shifts yet further in UI.

We focused our analyses on peak place cell activity, but future work could explore subtleties in firing rates (Sanders et al, 2019), and the relationship with theta rhythms (Chadwick et al, 2015). We only considered stable place cells before and after transitions; examining population turnover could yield further insights. To ensure robustness, we emphasized average spatial receptive fields, but tracking fast reward location changes remains essential. Finally, repeated switches, like those in UU, may eventually become expected, highlighting the need to understand how unknown unknowns transition to known unknowns in stochastic environments.

Future work should focus on deciphering the implementation processes underlying our findings. Plateau potentials generated by synchronized inputs from the entorhinal cortex and CA3 can lead to the formation of new feature-selective cells (Bittner et al, 2015). Furthermore, recent studies have highlighted enhanced reward-reference frame coding in the lateral entorhinal cortex (LEC) (Issa et al, 2024), and medial entorhinal cells are also attracted to goals (Boccara et al, 2019). Given that grid cells provide different spatial metrics and can anchor to task-relevant features (Peng et al, 2023), it would be natural to explore grid cell activity in the various conditions of our study. This might shed light on the structured diversity of CA1 place cells selectivity.

Task-relevant place cells selectivity could be driven by neuromodulatory inputs (Kaufman et al, 2020; Palacios-Filardo and Mellor, 2019; Palacios-Filardo et al, 2021). Evidence shows that acetylcholine, dopamine, noradrenaline and serotonin neuromodulatory systems provide signals associated with expectation, error and uncertainty, with their release reconfiguring hippocampal (and wider cortical) neuronal circuits to enable the update of estimates and memories (Dayan, 2012). Under this framework, the release of specific combinations of neuromodulators potentially codes for different types of uncertainty and could thereby influence the degree and type of place cell reorganisation. Indeed, dopaminergic and noradrenergic projections to CA1 from ventral tegmental area and locus coeruleus convey information about reward prediction errors (Cohen et al, 2012; Fiorillo et al, 2003; Schultz et al, 1997) and surprise (Fiorillo et al, 2003; Heer and Mark, 2023; Kaufman et al, 2020; McNamara et al, 2014) and can causally shape reward-related CA1 reorganisation (Kaufman et al, 2020; Krishnan et al, 2022), specifically in response to high reward prediction errors (Michon et al, 2021). Synaptic plasticity is the mechanism for place cell reorganisation and is regulated by neuromodulators in multiple ways (Palacios-Filardo and Mellor, 2019). For example, acetylcholine reprioritises entorhinal and CA3 inputs to CA1 reducing the internal representations from CA3 and enhancing external sensory input from entorhinal cortex (Hasselmo, 2006; Hasselmo and McGaughy, 2004; Palacios-Filardo et al, 2021) whilst also reconfiguring inhibitory networks (Haam et al, 2018; Leão et al, 2012) and enhancing dendritic excitability and synaptic plasticity (Buchanan et al, 2010; Dennis et al, 2016; Gu and Yakel, 2011; Teles-Grilo Ruivo and Mellor, 2013; Williams and Fletcher, 2019) in response to surprising events (Mineur et al, 2022; Ruivo et al, 2017). Thus, neuromodulators are an attractive mechanism linking detection of uncertainty to the update of hippocampal representations with new information.

In conclusion, we exploited the relative transparency of the spatial activity of hippocampal place cells in order to examine the effects of different forms of uncertainty about the location of reward, and, equally, used these different forms of uncertainty to enrich our understanding of the hippocampal code for space. Place cells exhibited impressive adaptation to the diverse statistical contingencies, with sub-populations adopting what we can see as different relevant reference frames. This sharpens the hippocampus’s role as not only a spatial navigator but also a flexible processor of uncertainty. By offering multiple reference frames depending on task-relevant features like reward, the hippocampus provides a robust framework for adapting to both expected and unexpected uncertainty. This flexibility suggests a novel mechanism by which the brain supports rapid decision-making under uncertainty —- crucial for survival in changing environments – and provides downstream circuits with a computationally sophisticated representation which can afford an attractive combination of specialization and generalization.

## Methods

### Mouse surgery

All experiments were approved and conducted in accordance with the Northwestern University Animal Care and Use Committee. Seven male P56-P63 mice (C57BL/6J, Jackson Laboratory, stock no.000664) were used in the experiments. To induce the expression of a calcium indicator, mice were first injected with AAV virus expressing jGCaMP8m (AAV1-syn-FLEX-jGCaMP8m-WPRE) (Zhang et al, 2023) into dorsal CA1 region of the right hippocampus (1.8mm lateral, 2.3mm caudal of Bregma, 1.25mm below the dura surface). After the injection, mice first recovered with ad libitum water for 1-2 days and then were subject to water restriction (0.8-1.2ml per day) until the end of all experiments. The weight of all mice was monitored and kept between 75%-80% of the original weight. After 3-5 days under water restriction, hippocampal cannula implant surgeries were performed above the injection site to allow optical access to dorsal CA1 of the hippocampus, as previously described (Dombeck et al, 2010). Briefly, cortex above the dorsal hippocampus was aspirated until the white matter of the external capsule was exposed. Phosphate buffer solution (PBS) was repeatedly applied until the bleeding stopped and a small drop of Kwik-Sil was applied to the tissue surface before the cannula was inserted. A head-plate and a ring were cemented on the skull using Meta-bond. Proper analgesic and anesthetic procedures were carried out according to the animal protocol. All mice were allowed to recover for 5-7 days before the start of behavioral training.

### Virtual reality and behavior task

Seven male mice were first separated into two groups, three and four mice for each group respectively. All mice were first habituated in the head-fixed VR setup (Sheffield et al, 2017) (with screen off) for one session (45 minutes), during which a couple of water rewards were delivered to the mice randomly to familiarize them with the lick port. Beginning from the second session, VR screens were turned on and both groups of mice were first trained in one visual environment to perform the URTask. Each training session lasted 45min to 1hr depending on how many laps the mice had run. Mice were considered well-trained if they satisfied both criteria: 1. They had to run at least 1 2 laps per minute; 2. They had to have anticipatory licking before the reward (anticipatory licking) for at least 50% of the laps; 3. Their behaviour is stable for three consecutive sessions, as measured by the average correlation coefficient of velocity and licking patterns across all laps. All mice reached this performance level after 8-10 session of training.

### Two-photon imaging

Two-photon calcium imaging of dorsal CA1 neurons was performed using a custom-built moveable objective microscope, with a 40x /0.8NA water immersion objective (LUMPlanFL N 3 40/0.8 W, Olympus), as described previously (Dombeck et al, 2010; Sheffield et al, 2017). The control software for two-photon scanning was ScanImage 5.1(Vidrio Technologies). Average laser power after the objective was around 60 100mW. Time-series movies of 12000 24000 frames, 512 x 256 pixels were acquired at 30Hz framerate. A Digidata1440A (Molecular Device) data acquisition system (Clampex 10.3) was used to record (at 1 kHz) and synchronize behavioral variables (licking, linear track position, velocity and reward delivery) with two-photon imaging frame time. During the same session, the imaging field stayed the same. During the consecutive imaging sessions, the imaging fields were not identical, although there might be overlap between the imaging fields.

### Image processing and ROI selection

Two-photon imaging time-series movies were first imported into Suite2p (Pachitariu et al, 2017) for rigid and non-rigid motion-correction. Putative cell (region of interest, ROIs) were extracted from motion-corrected movies using Suite2p.

**Table 1:** Suite2P Parameters

| Parameter | Value | Parameter | Value | Parameter | Value |
| --- | --- | --- | --- | --- | --- |
| nplanes | 1 | nchannels | 1 | functional_chan | 1 |
| tau | 0.6 | fs | 30 | do_bidiphase | 0 |
| bidiphase | 0 | multiplane_parallel | 0 | ignire_flyback | -1 |
| preclassify | 0 | save_mat | 1 | save_NWB | 0 |
| combined | 1 | reg_rig | 1 | reg_tif_chan2 | 0 |
| aspect | 1 | delete_bin | 0 | move_bin | 0 |
| do_registration | 1 | align_by_chan | 1 | nimg_init | 300 |
| batch_size | 500 | smooth_sigma | 1.15 | smooth_sigma_time | 0 |
| maxregshift | 0.1 | th_badframes | 1 | keep_movie_raw | 0 |
| two_step_registration | 0 | nonrigid | 1 | block_size | 32,64 |
| snr_thresh | 1.2 | maxregshiftNR | 5.0 | 1Preg | 0 |
| spatial_hp_reg | 32 | pre_smooth | 0 | spatial_taper | 40.0 |
| roidetect | 1 | denoise | 1 | spatial_scale | 0 |
| threshold_scaling | 2.0 | max_overlap | 0.75 | max_iterations | 20 |
| high_pass | 100.0 | spatial_hp_detect | 25 | anatomical_only | 0.0 |
| diameter | 0 |  |  |  |  |

Extracted ROI fluorescence traces were then exported from suite2P and imported into MATLAB for extracting significant calcium transients (Dombeck et al, 2010). For each ROI, the potential signal contamination from the surrounding neuropil was subtracted (after multiplied by a factor of 0.7) from the raw fluorescence signal. Slow time-course changes in the neuropil-corrected traces were removed by calculating the distribution of fluorescence in a 20-s time window around each time point and subtracting the 8th percentile value of the distribution. The baseline subtracted traces were then subjected to the analysis of the ratio of positive- to negative-going transients of various amplitudes and durations. This resulted in the identification of significant transients with less than 1% false positive rate. The significant transients were left untouched while all other values in the trace were set to 0. The resulting traces (referred to as ’changes in fluorescence’ in the following section) of all ROIs were used for further data analysis.

### Place cell spatial information test and identification

Fluorescence tuning maps were created by binning the position across the track into 60 bins and identifying the mean change in fluorescence when the animal was moving at least 0.1 cm per second. To test whether a cell is a place cell, we computed the spatial information (*I*) in bits per action potential for the fluorescence tuning map (Climer and Dombeck, 2021):

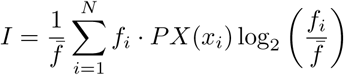

where *f̄* is the mean change in fluorescence, *N* is the number of bins, *f_i_* is the fluorescence change in the *i^th^* spatial bin, and *PX*(*x_i_*) is the probability that the animal is in the *i^th^* spatial bin. To build a null distribution of information, we circularly shuffled the fluorescence trace with a minimum shift of 15 seconds and recalculated the tuning map 1000 times. A cell was considered a significant place cell if it had higher information than 99% of these shuffled epochs, had an information value of at least 0.5 bits per action potential.

### Trial inclusion criteria

The position of reward consumption was defined as the first lick after reward delivery on every trial. As animals were engaged in the task, on most trials, licking was very close to reward delivery. The reward zone was then defined as the zone between the most proximal reward consumption position, until the most distal reward consumption position.

In order to obtain a meaningful reward zone, we excluded 2.5% of the trials (35 out of 1376 total trials included in this paper) that were outlier in the distance at which the reward was consumed after delivery. This selection criteria generated a threshold of approximately 11 cm between reward delivery and reward consumption, therefore excluding trials in which the reward was not consumed, or was consumed after this distance. Supplementary Figure **??** shows the histogram of consumption distance from reward delivery, which we also consider as a marker for engagement in the task.

### Trial separation

We separated proximal, middle and distal rewards by dividing the reward zone in 3 bins of identical length. The trials in which the reward was consumed in the first (resp. second, third) bin were labelled ’proximal’ (resp. ’middle’, ’distal’).

### Behaviour analysis

We excluded from all analyses the teleportation phase (during which the screen went black), and all datapoint at which the velocity fell under 0.1 cm/s.

Analyses were performed using custom Python code. To calculate the lick rate and velocity patterns (figures 2a;b, figure 3a;b), we averaged the lick rate and velocity trace, downsampled at 30 Hz, over a position vector covering all position values (from 0 to 3m) with a bin size of 10 cm. To compute averages, we extracted the values of the behavioral variables for the cases in which the position trace was within each position bin, and computed averages weighted by the time spent in each position bin. For figure 2a, for every session average-value, we computed the average over all trials for LU and divided it by the maximum value over the session. We then averaged this value across sessions and animals. For figure 2b, for EU, we computed the average on proximal, middle and distal trials, and normalised it to the maximum value of the average computed over the full session. We then averaged these values across sessions and animals.

### Place cell activity analysis

For all place cells analyses, we excluded periods in which the animal ran with a velocity less than 1cm/s, and the teleportation corridor. For figure 2d;e, Figure 3c;d, and Figure 3b, each place cell’s activity was averaged similarly to behavioural variables: the average place cell activity over the session was computed by averaging the activity per position bin across every trial weighted by the time spent in each position bin. Place maps in figures 2d;e show the average activity of cells on odd trials, ordered based on the location of the peak activity on even trials. Place map plots were produced by normalizing the average activity of every cell on odd trials by its maximum value.

For switch sessions (place maps in figures 3c;d), place map plots before the switch were produced by normalising the average activity of every cell on all trials before the switch by its maximum value. Similarly, place map plots after the switch were produced by normalising the average activity of every cell on all trials after the switch by its maximum value.

### Peak activity analysis

The position of maximum activity was extracted as the location of the 10cm bin in which the average activity of the cell was greatest. For figure 2e, we considered the average activity on proximal and distal groups of trials. For figure 3e, the average was computed over trials before (x-axis) and after (y-axis) the reward switch separately.

For figures 2f and 3e, the x and y coordinates are fitted with gaussian kde function from the scipy.stats module, which estimates the probability density function (PDF) of a random variable in a non-parametric way. The heatmap shows this Gaussian fitted density estimation.

### Reward and warped reference frame

The reward reference frame was obtained by computing positions relative to the position of the consumption of the reward at every trial, and using 10cm position bins.

The warped reference frame was obtained by creating a warped vector interpolating the position in 20 bins between the start of the track and the reward location, and 20 bins from the reward location to the end of the track at every trial. These new bins were then the basis for all averages.

### Place cell identification

In figure 2g,’Position-stable” cells were place cells that passed the place cell test and which position of peak of activity on proximal and distal trials were at most 15cm apart.

In figure 3f,’Position-stable” cells were place cells that passed the place cell test before and after the switch and which position of peak of activity before and after the switch were at most 15cm apart.

In figure 3g, ’reward-peaking’ cells were place cells that passed the place cell test before and after the switch and whose positions of peak of activity in the reward reference frame before and after the switch were between -15 and +20 cm.

In figure 3h, ’Warped’ place cells were place cells that passed the place cell test before and after the switch and which position of peak of activity in the warped reference frame before and after the switch were identical with + or - 3 warped units, and which position of maximum activity followed the reward.

### Cell percentage and cell percentage per cm Statistical analyses

All statistics were done using the package ’statsmodels’ in python.

To compare percentages, we used the percentage z-test, and for 1-sided proportion z-test to test for directionality. To compare distributions, we used the Kolmogorov-Smirnov test.

## Data availability

The data will be made freely available following publication.

## Code availability

All computer programs will be made freely available following publication.

## Acknowledgements

We are grateful to Claudia Clopath, Matt Jones, Zach Mainen, Tony Pickering and Mark Walton for influential discussions about the design of the task. We thank Marielena Sosa, Mark Plitt and Lisa Giocomo for sharing their results prior to publication.

## Funding

Funding was from the Max Planck Society (CT, PD) and the Humboldt Foundation (PD). PD is a member of the Machine Learning Cluster of Excellence, EXC number 2064/1 – Project number 39072764 and of the Else Kröner Medical Scientist Kolleg “ClinbrAIn: Artificial Intelligence for Clinical Brain Research”. Funding for JRM from Wellcome Trust (101029/Z/13/Z) and Biotechnology and Biological Sciences Research Council (BBSRC, BB/V001728/1, BB/N013956/1). FX was supported by National Institute of Mental Health Training Program in Neurobiology of Information Storage (T32MH067564) predoctoral fellowship.

## Author Credit Contribution

**Fig. 5:**
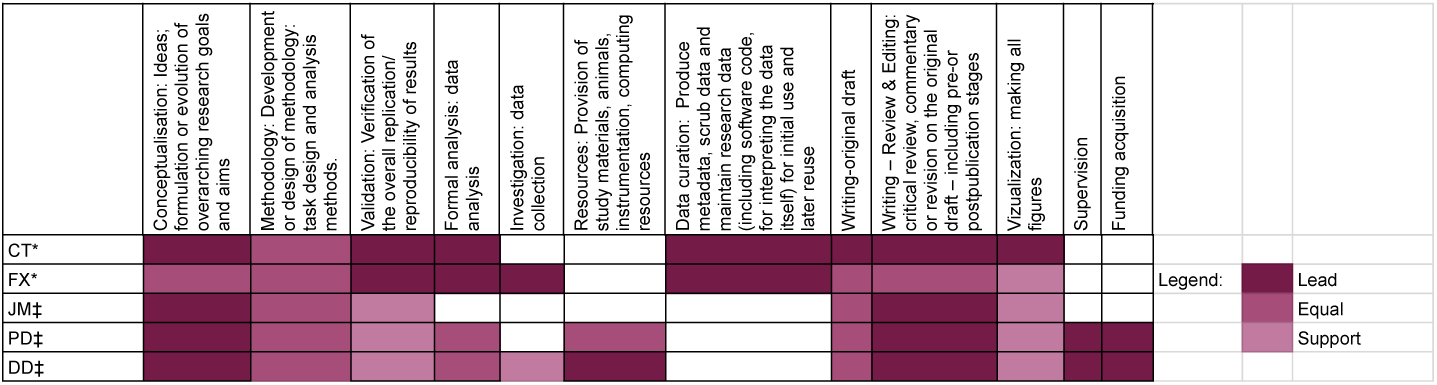
CRediT. CRediT contribution matrix. Color code refers to the level of contribution per category, as previously used (Tay, 2021). Categories reflect the ones published in the original CRediT taxonomy in (Brand et al, 2015).

## Conflict of interest/Competing interests

The authors declare no conflict of interest.

## Notes

### Competing Interest Statement

The authors have declared no competing interest.

### Summary of Updates

Added a contributor to data collection as author.

## References

1. Anderson MI, Jeffery KJ (2003) Heterogeneous modulation of place cell firing by changes in context. Journal of Neuroscience 23(26):8827–8835

2. Aoki Y, Igata H, Ikegaya Y, et al (2019) The integration of goal-directed signals onto spatial maps of hippocampal place cells. Cell reports 27(5):1516–1527

3. Bast T, Wilson IA, Witter MP, et al (2009) From rapid place learning to behavioral performance: a key role for the intermediate hippocampus. PLoS biology 7(4):e1000089

4. Behrens TE, Woolrich MW, Walton ME, et al (2007) Learning the value of information in an uncertain world. Nature neuroscience 10(9):1214–1221

5. Best PJ, White AM, Minai A (2001) Spatial processing in the brain: the activity of hippocampal place cells. Annual review of neuroscience 24(1):459–486

6. Bittner KC, Grienberger C, Vaidya SP, et al (2015) Conjunctive input processing drives feature selectivity in hippocampal ca1 neurons. Nature neuroscience 18(8):1133–1142

7. Boccara CN, Nardin M, Stella F, et al (2019) The entorhinal cognitive map is attracted to goals. Science 363(6434):1443–1447

8. Bostock E, Muller RU, Kubie JL (1991) Experience-dependent modifications of hippocampal place cell firing. Hippocampus 1(2):193–205

9. Brand A, Allen L, Altman M, et al (2015) Beyond authorship: Attribution, contribution, collaboration, and credit. Learned Publishing 28(2)

10. Buchanan KA, Petrovic MM, Chamberlain SE, et al (2010) Facilitation of long-term potentiation by muscarinic m1 receptors is mediated by inhibition of sk channels. Neuron 68(5):948–963

11. Burgess N, O’Keefe J (1996) Neuronal computations underlying the firing of place cells and their role in navigation. Hippocampus 6(6):749–762

12. Burgess N, Recce M, O’Keefe J (1995) Hippocampus: spatial models. The handbook of brain theory and neural networks pp 468–472

13. Chadwick A, van Rossum MC, Nolan MF (2015) Independent theta phase coding accounts for ca1 population sequences and enables flexible remapping. Elife 4:e03542

14. Climer JR, Dombeck DA (2021) Information Theoretic Approaches to Deciphering the Neural Code with Functional Fluorescence Imaging. eNeuro 8(5)

15. Cohen JY, Haesler S, Vong L, et al (2012) Neuron-type-specific signals for reward and punishment in the ventral tegmental area. nature 482(7383):85–88

16. Cohen JY, Amoroso MW, Uchida N (2015) Serotonergic neurons signal reward and punishment on multiple timescales. Elife 4:e06346

17. Dayan P (2012) Twenty-five lessons from computational neuromodulation. Neuron 76(1):240–256

18. Dennis SH, Pasqui F, Colvin EM, et al (2016) Activation of muscarinic m1 acetylcholine receptors induces long-term potentiation in the hippocampus. Cerebral cortex 26(1):414–426

19. Dombeck DA, Harvey CD, Tian L, et al (2010) Functional imaging of hippocampal place cells at cellular resolution during virtual navigation. Nature neuroscience 13(11):1433–1440

20. Dupret D, O’Neill J, Pleydell-Bouverie B, et al (2010) The reorganization and reactivation of hippocampal maps predict spatial memory performance. Nature neuroscience 13(8):995–1002

21. Fiorillo CD, Tobler PN, Schultz W (2003) Discrete coding of reward probability and uncertainty by dopamine neurons. Science 299(5614):1898–1902

22. Foster DJ, Morris RG, Dayan P (2000) A model of hippocampally dependent navigation, using the temporal difference learning rule. Hippocampus 10(1):1–16

23. Frank LM, Stanley GB, Brown EN (2004) Hippocampal plasticity across multiple days of exposure to novel environments. Journal of Neuroscience 24(35):7681–7689

24. Gauthier JL, Tank DW (2018) A dedicated population for reward coding in the hippocampus. Neuron 99(1):179–193

25. Gothard KM, Skaggs WE, McNaughton BL (1996) Dynamics of mismatch correction in the hippocampal ensemble code for space: interaction between path integration and environmental cues. Journal of Neuroscience 16(24):8027–8040

26. Gu Z, Yakel JL (2011) Timing-dependent septal cholinergic induction of dynamic hippocampal synaptic plasticity. Neuron 71(1):155–165

27. Haam J, Zhou J, Cui G, et al (2018) Septal cholinergic neurons gate hippocampal output to entorhinal cortex via oriens lacunosum moleculare interneurons. Proceedings of the National Academy of Sciences 115(8):E1886–E1895

28. Hasselmo ME (2006) The role of acetylcholine in learning and memory. Current opinion in neurobiology 16(6):710–715

29. Hasselmo ME, McGaughy J (2004) High acetylcholine levels set circuit dynamics for attention and encoding and low acetylcholine levels set dynamics for consolidation. Progress in brain research 145:207–231

30. Heer CM, Mark E (2023) Distinct catecholaminergic pathways projecting to hippocampal ca1 transmit contrasting signals during behavior and learning. bioRxiv

31. Hill A (1978) First occurrence of hippocampal spatial firing in a new environment. Experimental neurology 62(2):282–297

32. Hollup SA, Molden S, Donnett JG, et al (2001) Accumulation of hippocampal place fields at the goal location in an annular watermaze task. Journal of Neuroscience 21(5):1635–1644

33. Hsu M, Bhatt M, Adolphs R, et al (2005) Neural systems responding to degrees of uncertainty in human decision-making. Science 310(5754):1680–1683

34. Hüllermeier E, Waegeman W (2021) Aleatoric and epistemic uncertainty in machine learning: An introduction to concepts and methods. Machine learning 110(3):457–506

35. Issa JB, Radvansky BA, Xuan F, et al (2024) Lateral entorhinal cortex subpopulations represent experiential epochs surrounding reward. Nature neuroscience 27(3):536–546

36. Jarzebowski P, Hay YA, Grewe BF, et al (2022) Different encoding of reward location in dorsal and intermediate hippocampus. Current Biology 32(4):834–841

37. Kaufman AM, Geiller T, Losonczy A (2020) A role for the locus coeruleus in hippocampal ca1 place cell reorganization during spatial reward learning. Neuron 105(6):1018–1026

38. Kentros C, Hargreaves E, Hawkins RD, et al (1998) Abolition of long-term stability of new hippocampal place cell maps by nmda receptor blockade. Science 280(5372):2121–2126

39. Kleinknecht KR, Bedenk BT, Kaltwasser SF, et al (2012) Hippocampus-dependent place learning enables spatial flexibility in c57bl6/n mice. Frontiers in behavioral neuroscience 6:87

40. Krishnan S, Heer C, Cherian C, et al (2022) Reward expectation extinction restructures and degrades ca1 spatial maps through loss of a dopaminergic reward proximity signal. Nature communications 13(1):6662

41. Kumaran D, Maguire EA (2007) Which computational mechanisms operate in the hippocampus during novelty detection? Hippocampus 17(9):735–748

42. Leão RN, Mikulovic S, Leão KE, et al (2012) Olm interneurons differentially modulate ca3 and entorhinal inputs to hippocampal ca1 neurons. Nature neuroscience 15(11):1524–1530

43. Leutgeb JK, Leutgeb S, Treves A, et al (2005a) Progressive transformation of hippocampal neuronal representations in “morphed” environments. Neuron 48(2):345–358

44. Leutgeb S, Leutgeb JK, Barnes CA, et al (2005b) Independent codes for spatial and episodic memory in hippocampal neuronal ensembles. Science 309(5734):619–623

45. Markus EJ, Qin YL, Leonard B, et al (1995) Interactions between location and task affect the spatial and directional firing of hippocampal neurons. Journal of Neuroscience 15(11):7079–7094

46. McGuire JT, Nassar MR, Gold JI, et al (2014) Functionally dissociable influences on learning rate in a dynamic environment. Neuron 84(4):870–881

47. McKenzie S, Robinson NT, Herrera L, et al (2013) Learning causes reorganization of neuronal firing patterns to represent related experiences within a hippocampal schema. Journal of Neuroscience 33(25):10243–10256

48. McKenzie S, Frank AJ, Kinsky NR, et al (2014) Hippocampal representation of related and opposing memories develop within distinct, hierarchically organized neural schemas. Neuron 83(1):202–215

49. McNamara CG, Tejero-Cantero Á, Trouche S, et al (2014) Dopaminergic neurons promote hippocampal reactivation and spatial memory persistence. Nature neuroscience 17(12):1658–1660

50. McNaughton N, Bannerman D (2024) The homogenous hippocampus: How hippocampal cells process available and potential goals. Progress in neurobiology p 102653

51. Michon F, Krul E, Sun JJ, et al (2021) Single-trial dynamics of hippocampal spatial representations are modulated by reward value. Current Biology 31(20):4423–4435

52. Mineur YS, Mose TN, Vanopdenbosch L, et al (2022) Hippocampal acetylcholine modulates stress-related behaviors independent of specific cholinergic inputs. Molecular psychiatry 27(3):1829–1838

53. Morris RGM, Davis S, Butcher S (1990) Hippocampal synaptic plasticity and nmda receptors: a role in information storage? Philosophical Transactions of the Royal Society of London Series B: Biological Sciences 329(1253):187–204

54. Moser EI, Kropff E, Moser MB (2008) Place cells, grid cells, and the brain’s spatial representation system. Annu Rev Neurosci 31(1):69–89

55. Muller R (1996) A quarter of a century of place cells. Neuron 17(5):813–822

56. Muller RU, Kubie JL (1987) The effects of changes in the environment on the spatial firing of hippocampal complex-spike cells. Journal of Neuroscience 7(7):1951–1968

57. Muzzio IA, Kentros C, Kandel E (2009) What is remembered? role of attention on the encoding and retrieval of hippocampal representations. The Journal of physiology 587(12):2837–2854

58. Nassar MR, McGuire JT, Ritz H, et al (2019) Dissociable forms of uncertainty-driven representational change across the human brain. Journal of Neuroscience 39(9):1688–1698

59. Nitz D (2009) Parietal cortex, navigation, and the construction of arbitrary reference frames for spatial information. Neurobiology of learning and memory 91(2):179–185

60. O’Keefe J, Dostrovsky J (1971) The hippocampus as a spatial map: preliminary evidence from unit activity in the freely-moving rat. Brain research

61. Pachitariu M, Stringer C, Dipoppa M, et al (2017) Suite2p: beyond 10,000 neurons with standard two-photon microscopy. bioRxiv

62. Palacios-Filardo J, Mellor JR (2019) Neuromodulation of hippocampal long-term synaptic plasticity. Current opinion in neurobiology 54:37–43

63. Palacios-Filardo J, Udakis M, Brown GA, et al (2021) Acetylcholine prioritises direct synaptic inputs from entorhinal cortex to ca1 by differential modulation of feedforward inhibitory circuits. Nature communications 12(1):5475

64. Peng JJ, Throm B, Najafian Jazi M, et al (2023) Grid cells perform path integration in multiple reference frames during self-motion-based navigation. bioRxiv pp 2023–12

65. Plitt MH, Giocomo LM (2021) Experience-dependent contextual codes in the hippocampus. Nature neuroscience 24(5):705–714

66. Preuschoff K, ’t Hart BM, Einhäuser W (2011) Pupil dilation signals surprise: Evidence for nora-drenaline’s role in decision making. Frontiers in neuroscience 5:115

67. Radvansky BA, Oh JY, Climer JR, et al (2021) Behavior determines the hippocampal spatial mapping of a multisensory environment. Cell reports 36(5)

68. Ruivo LMTG, Baker KL, Conway MW, et al (2017) Coordinated acetylcholine release in prefrontal cortex and hippocampus is associated with arousal and reward on distinct timescales. Cell reports 18(4):905–917

69. Sanders H, Ji D, Sasaki T, et al (2019) Temporal coding and rate remapping: Representation of nonspatial information in the hippocampus. Hippocampus 29(2):111–127

70. Sanders H, Wilson MA, Gershman SJ (2020) Hippocampal remapping as hidden state inference. Elife 9:e51140

71. Sarel A, Finkelstein A, Las L, et al (2017) Vectorial representation of spatial goals in the hippocampus of bats. Science 355(6321):176–180

72. Sato M, Mizuta K, Islam T, et al (2020) Distinct mechanisms of over-representation of landmarks and rewards in the hippocampus. Cell reports 32(1)

73. Schultz W, Dayan P, Montague PR (1997) A neural substrate of prediction and reward. Science 275(5306):1593–1599

74. Sheffield ME, Dombeck DA (2019) Dendritic mechanisms of hippocampal place field formation. Current opinion in neurobiology 54:1–11

75. Sheffield ME, Adoff MD, Dombeck DA (2017) Increased prevalence of calcium transients across the dendritic arbor during place field formation. Neuron 96(2):490–504

76. Skaggs WE, McNaughton BL (1998) Spatial firing properties of hippocampal ca1 populations in an environment containing two visually identical regions. Journal of Neuroscience 18(20):8455–8466

77. Soltani A, Izquierdo A (2019) Adaptive learning under expected and unexpected uncertainty. Nature Reviews Neuroscience 20(10):635–644

78. Sosa M, Giocomo LM (2021) Navigating for reward. Nature Reviews Neuroscience 22(8):472–487

79. Sosa M, Plitt MH, Giocomo LM (2023) Hippocampal sequences span experience relative to rewards. bioRxiv

80. Steele R, Morris R (1999) Delay-dependent impairment of a matching-to-place task with chronic and intrahippocampal infusion of the nmda-antagonist d-ap5. Hippocampus 9(2):118–136

81. Sun W, Winnubst J, Natrajan M, et al (2023) Learning produces a hippocampal cognitive map in the form of an orthogonalized state machine. bioRxiv pp 2023–08

82. Tanni S, De Cothi W, Barry C (2022) State transitions in the statistically stable place cell population correspond to rate of perceptual change. Current Biology 32(16):3505–3514

83. Tay A (2021) Researchers are embracing visual tools to give fair credit for work on papers. Nature Index vom 22:2021

84. Teles-Grilo Ruivo LM, Mellor JR (2013) Cholinergic modulation of hippocampal network function. Frontiers in synaptic neuroscience 5:2

85. Tessereau C, O’Dea R, Coombes S, et al (2021) Reinforcement learning approaches to hippocampus-dependent flexible spatial navigation. Brain and Neuroscience Advances 5:2398212820975634

86. Vaidya SP, Chitwood RA, Magee JC (2023) The formation of an expanding memory representation in the hippocampus. biorxiv pp 2023–02

87. Vinogradova OS (2001) Hippocampus as comparator: role of the two input and two output systems of the hippocampus in selection and registration of information. Hippocampus 11(5):578–598

88. Wallenstein GV, Hasselmo ME, Eichenbaum H (1998) The hippocampus as an associator of discontiguous events. Trends in neurosciences 21(8):317–323

89. Williams SR, Fletcher LN (2019) A dendritic substrate for the cholinergic control of neocortical output neurons. Neuron 101(3):486–499

90. Wills TJ, Lever C, Cacucci F, et al (2005) Attractor dynamics in the hippocampal representation of the local environment. Science 308(5723):873–876

91. Wood ER, Dudchenko PA, Robitsek RJ, et al (2000) Hippocampal neurons encode information about different types of memory episodes occurring in the same location. Neuron 27(3):623–633

92. Yu AJ, Dayan P (2005) Uncertainty, neuromodulation, and attention. Neuron 46(4):681–692

93. Zeithamova D, Gelman BD, Frank L, et al (2018) Abstract representation of prospective reward in the hippocampus. Journal of Neuroscience 38(47):10093–10101

94. Zhang Y, Ŕozsa M, Liang Y, et al (2023) Fast and sensitive gcamp calcium indicators for imaging neural populations. Nature 615(7954):884–891

95. Zinyuk L, Kubik S, Kaminsky Y, et al (2000) Understanding hippocampal activity by using purposeful behavior: place navigation induces place cell discharge in both task-relevant and task-irrelevant spatial reference frames. Proceedings of the National Academy of Sciences 97(7):3771–3776

