## Supplementary material for "Navigating uncertainty: reward location variability induces reorganization of hippocampal spatial representations"

### Code availability

All computer programs will be made freely available following publication.

### Supplementary material

Please see supplementary figures.

**Fig. S1**: **Supplementary Figure 1: Reward consumption statistics and trial inclusion criteria**

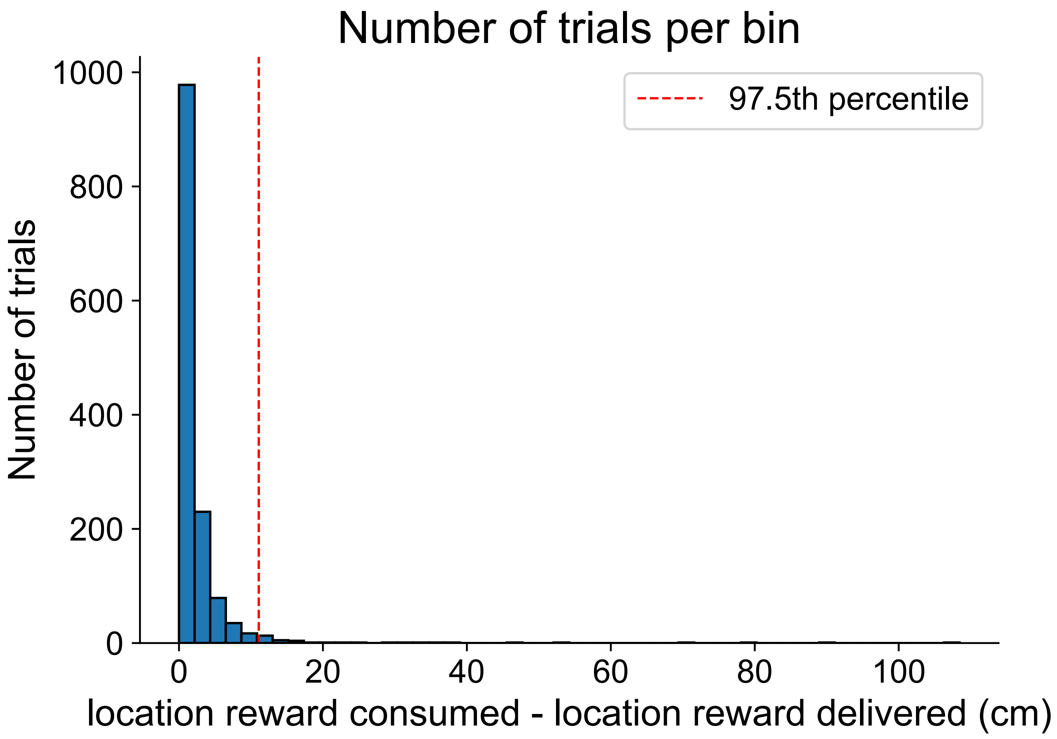


Number of trials (y-axis) per bin of values of distance between reward consumption and reward delivery (x-axis). Red vertical bar shows the 97.5% criteria used for inclusion in analyses shown here.

---

**Fig. S2**: **Supplementary Figure 2a: Additional place map analyses and individual place maps.**

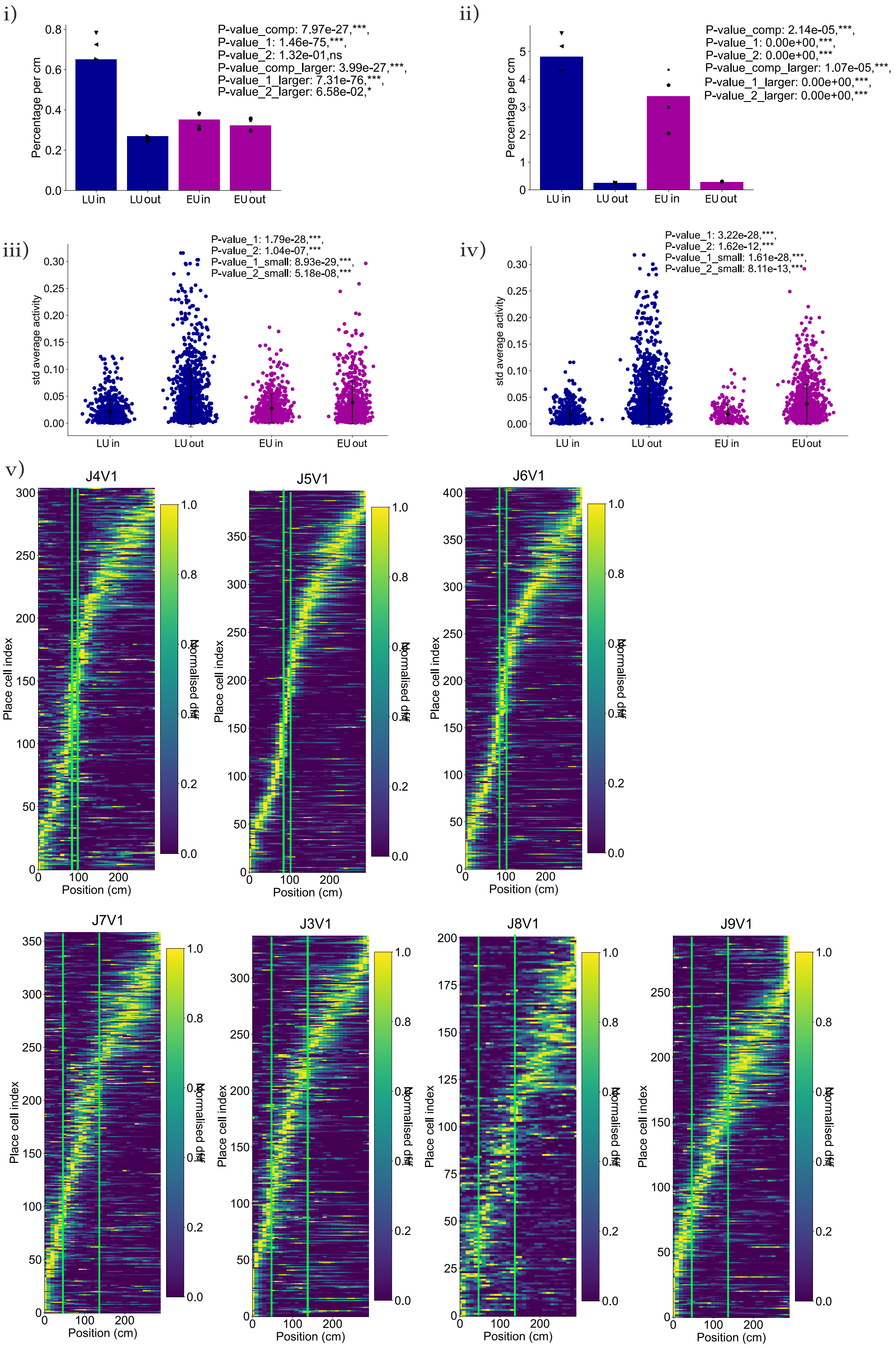


i: Density of place cell per cm at the reward zone (±15cm - "in") and out of the reward zone ("out") for LU (darkblue) and EU (darkpink). Symbols show single session percentages per cm. Comparisons: LU comparison in/out proportion z-test: $p = 1.5 \times 10^{-75}$, LU comparison in>out 1-sided proportion z-test:

$p = 7.3 \times 10^{-76}$, EU comparison in/out proportion z-test: $p = 1.3 \times 10^{-1}$, EU comparison in>out 1-sided proportion z-test: $p = 6.6 \times 10^{-2}$, comparison z-test LU-in/EU-in $p = 8 \times 10^{-27}$, 1-sided comparison z-test LU-in>EU-in $p = 4 \times 10^{-27}$.
ii: Similar to c:i) but in a reward-reference frame.
Comparisons: LU comparison in/out proportion z-test: $p < 2.2 \times 10^{-308}$, LU comparison in>out 1-sided proportion z-test: $p < 2.2 \times 10^{-308}$, EU comparison in/out proportion z-test: $p < 2.2 \times 10^{-308}$, EU comparison in>out 1-sided proportion z-test: $p < 2.2 \times 10^{-308}$, comparison z-test LU-in/EU-in $p = 2.14 \times 10^{-5}$, 1-sided comparison z-test LU-in>EU-in $p = 1.07 \times 10^{-5}$.
iii: Standard deviation of the average activity of cells. Each dot shows the spatial std of the average activity over the whole session for a single cell. Points are jittered according to their std values for density visualisation, for cells in the reward zone (”in) and out of the reward zone (”out”), for LU (blue) and EU (pink).
iv: Similar to iii: but in a reward reference frame.
v: Place maps as in 2 for each animal.
Green thick lines show the reward zone.

**Fig. S3**: **Supplementary Figure 2b: Velocity and licking patterns for individual animal.**

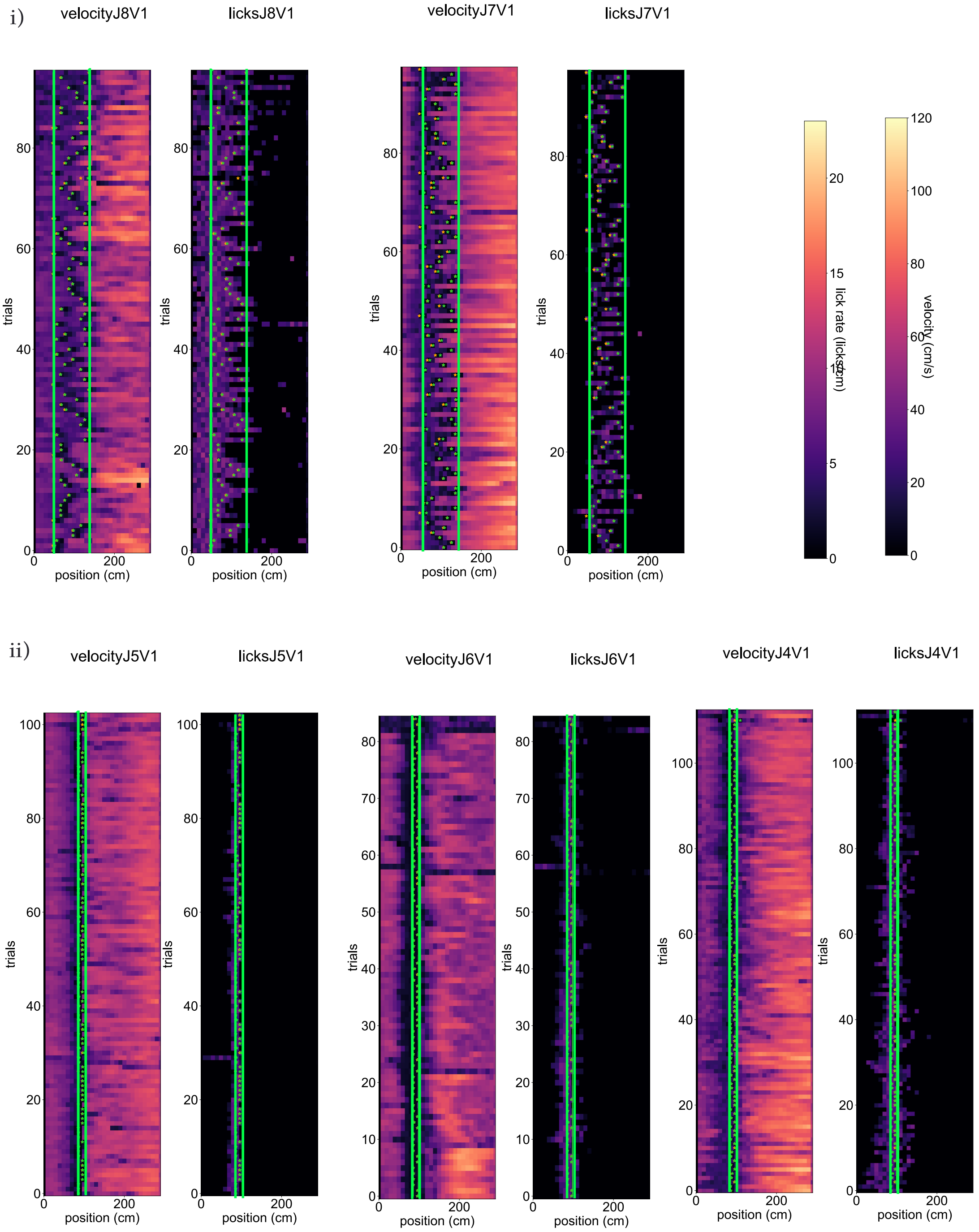


i: Velocity and licking pattern for each animal in EU.
ii: Similar to i: in EU.
For i and ii, the colorbars giving value ranges for licking and velocity are shown as inserts.
Green thick lines show the reward zone, green stars the reward consumption locations, and the orange stars the reward delivery location.

---

**Fig. S4: Supplementary Figure 2c: Individual cell examples.**

i)

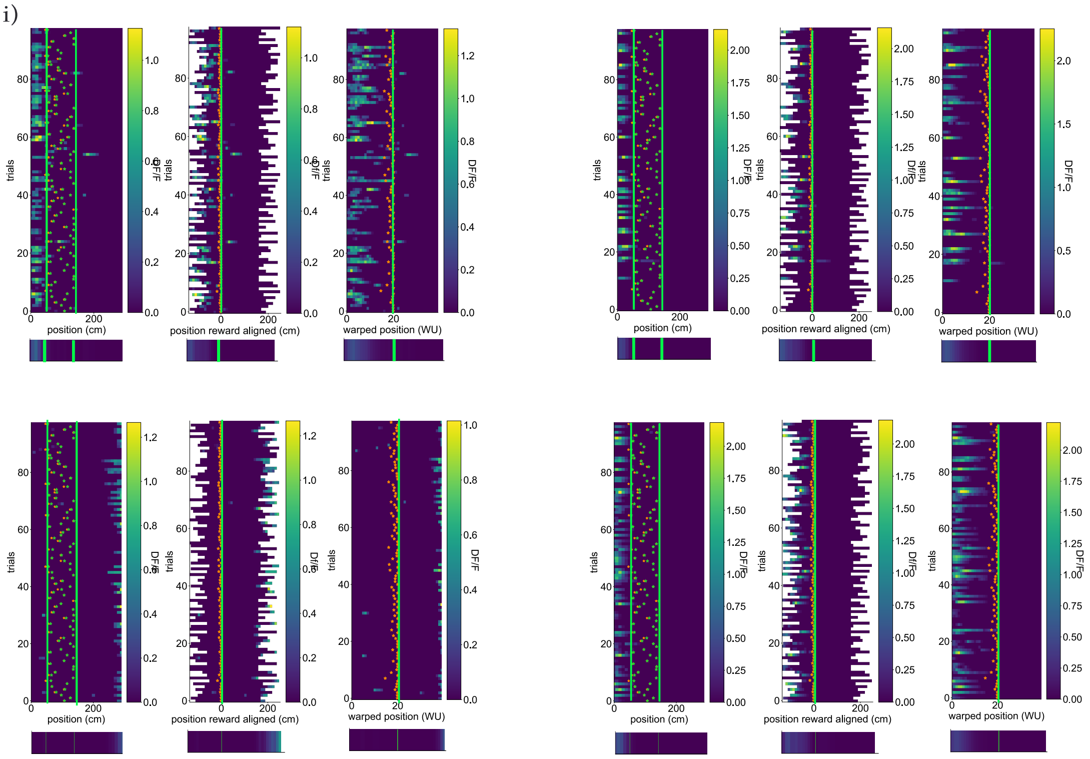


ii)

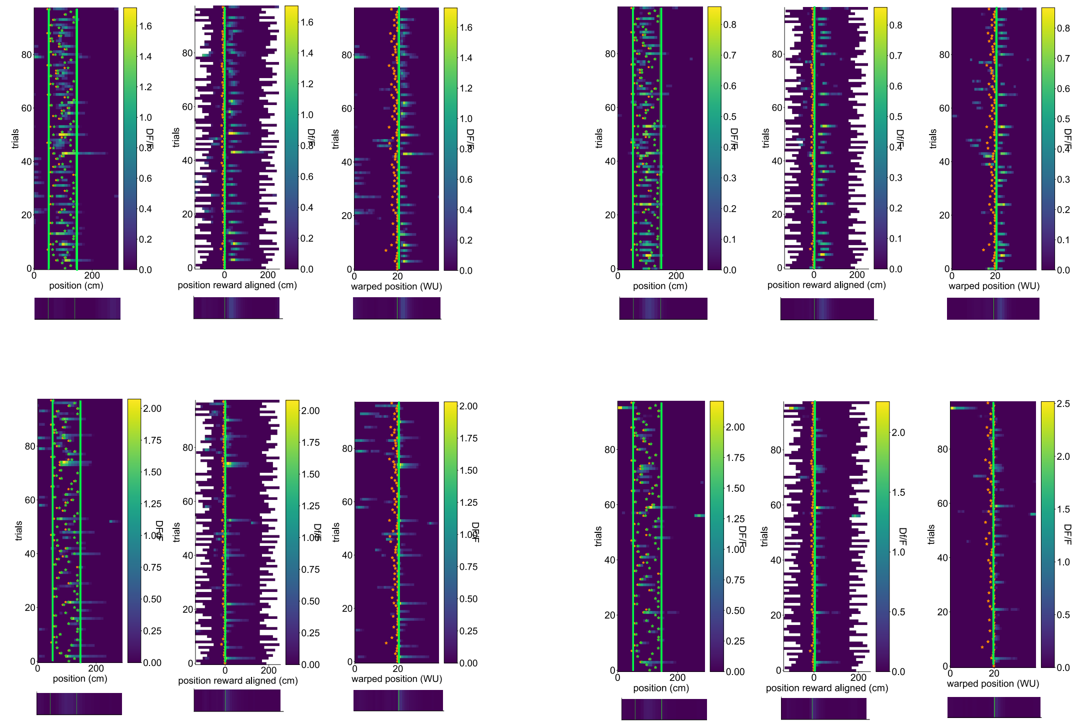

iii)

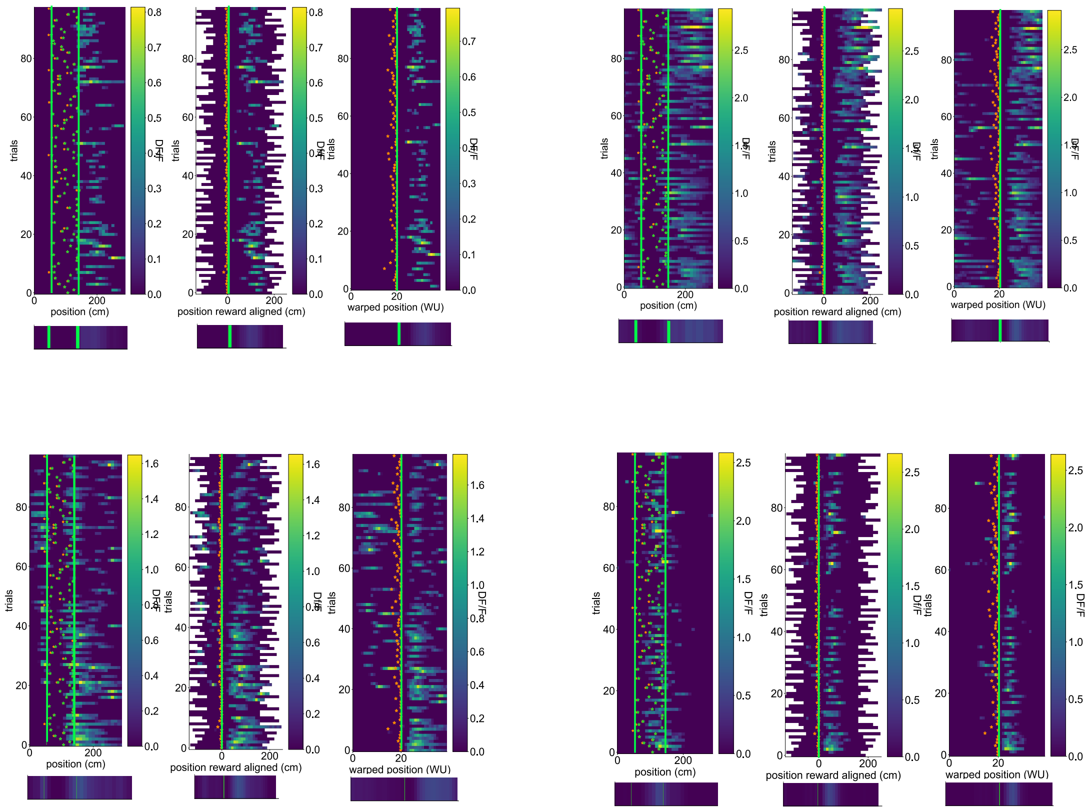


i: Four position-stable cells: example cells that keep their maximum of activity between proximal and distal reward trials in position reference frame. For each cell, the activity is shown in a position reference frame (left), in a reward reference frame (middle), in a warped reference frame (right).
ii: Similar to i: for four reward-stable cells: cells that keep their maximum of activity between proximal and distal reward trials in a reward reference frame.
iii: Similar to (i,ii) for four warped-stable cells: cells that keep their maximum of activity between proximal and distal reward trials in a warped reference frame.
Green thick lines show the reward zone, green stars the reward consumption locations, and the orange stars the reward delivery location.

---

**Fig. S5**: **Supplementary Figure 3a: Additional place map analyses.**

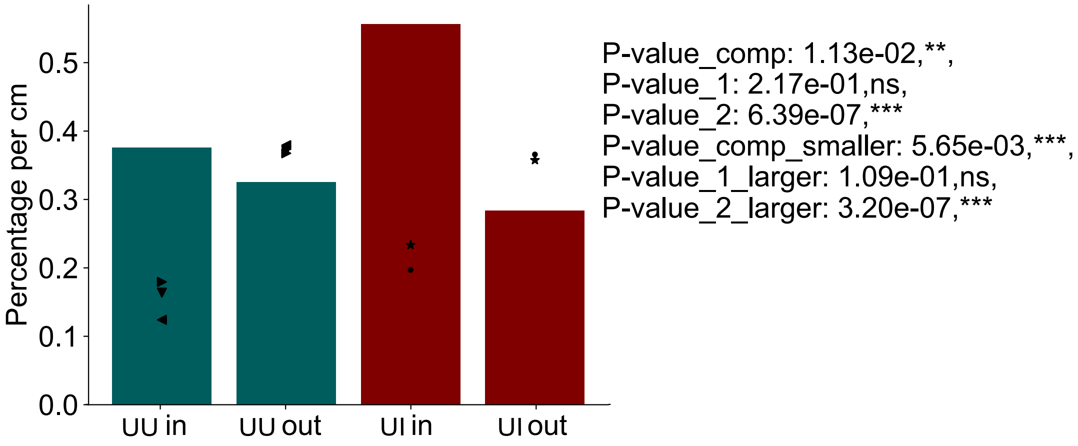

Density of cells per cm at the reward zone ("in") and outside the reward zone ("out") after the switch for UU (green) and UI (red). Symbols show single session percentages per cm. Comparisons: UU comparison in/out proportion z-test: $p = 2.17 \times 10^{-1}$, UU comparison in>out 1-sided proportion z-test: $p < 1 \times 10^{-1}$, UI comparison in/out proportion z-test: $p = 6.4 \times 10^{-7}$, UI comparison in>out 1-sided proportion z-test: $p = 3.2 \times 10^{-7}$, comparison z-test UU-in/UI-in $p = 1.13 \times 10^{-2}$, 1-sided comparison z-test UU-in<UI-in $p = 5.7 \times 10^{-3}$.

---

**Fig. S6**: **Supplementary Figure 3b: Reproduction of results for animal J7 only.**

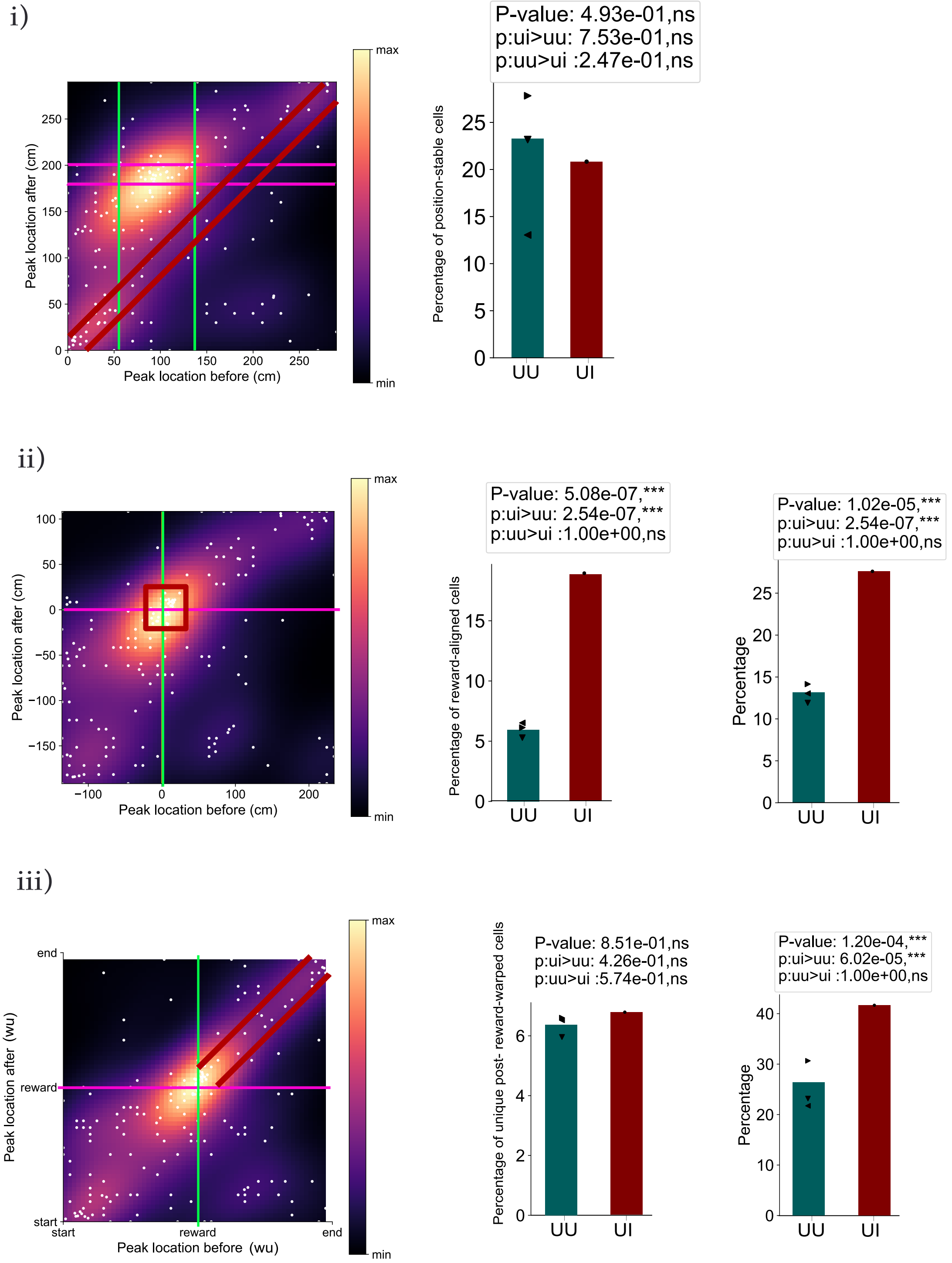


Similar to 3e if excluding animal J8 and including only J7 in the UI data.
i: in a position reference frame. Left shows scatter plot as in 3e:i, middle the diagonal count as in 3f.
ii: Similar to i: in a reward reference frame. Middle shows the count of cells stably following the reward before and after the switch, excluding any position-aligned cells, as in 3g, right shows the full diagonal count, as in 4c:ii.

iii: Similar to (i,ii) in a warped reference frame. Middle shows the post-reward diagonal count, excluding any reward nor position stable-cells, as in 3h. Right shows the warped diagonal count, as in 4d:i.

---

**Fig. S7**: **Supplementary Figure 3c: Reproduction of results for animal J7 only.**

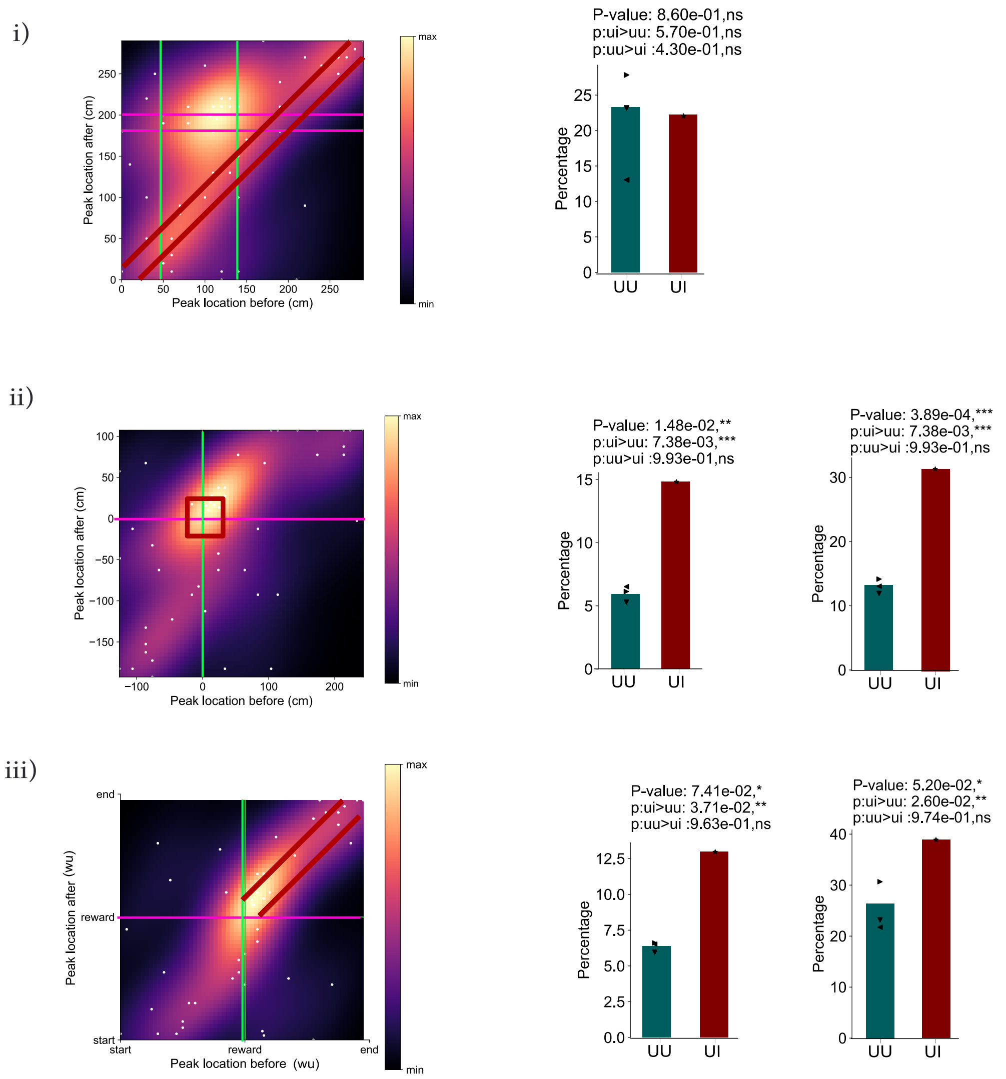


Similar to S6, but keeping only J8 and discarding J7.

**Fig. S8**: **Supplementary Figure 3d: Lick rate and velocity patterns for individual animal.**

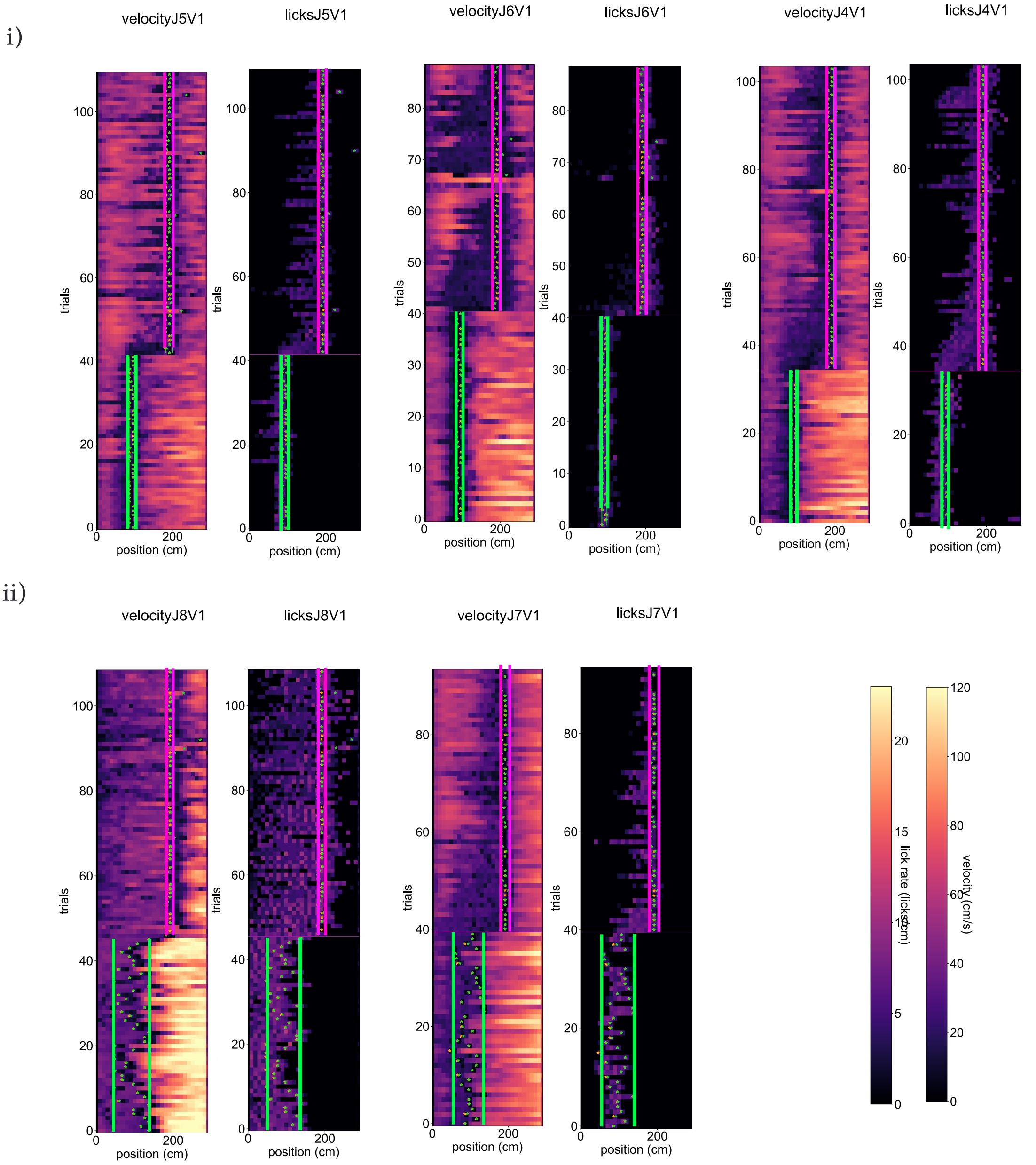


i: Velocity and licking pattern for each animal in UU.
ii: Similar to i: in UI.
For i and ii, the colorbars giving value ranges for licking and velocity are shown as inserts.
Green thick lines show the full reward zone before the switch, pink lines the reward zone after the switch, green stars the reward consumption locations, and the orange stars the reward delivery location.

---

**Fig. S9**: **Supplementary Figure 3e: Individual place maps.**

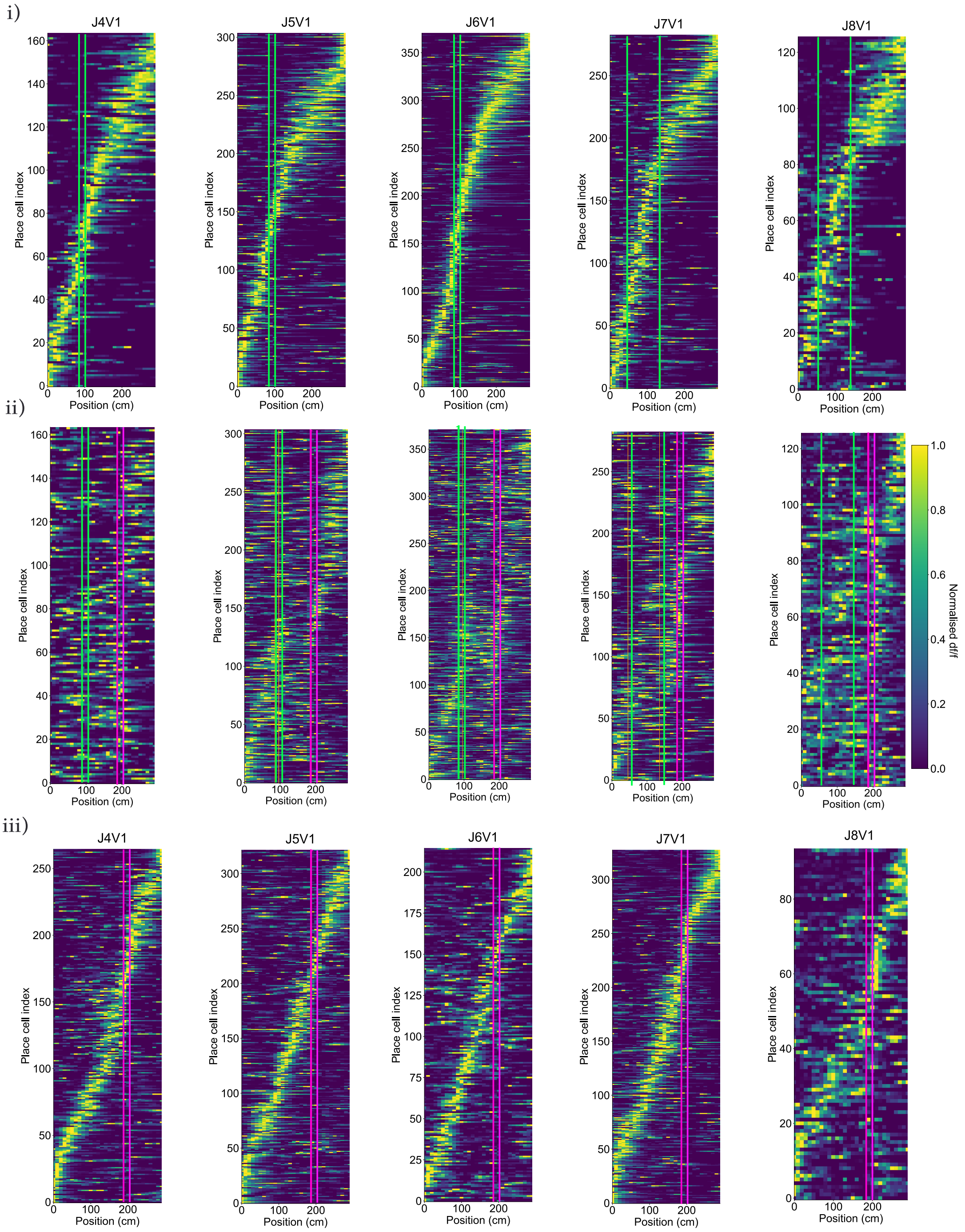


i: Individual normalized place maps before the switch.
ii: Individual normalized place maps after the switch, in the same order than i:
iii: Individual new normalized place maps after the switch.
Green thick lines show the full reward zone before the switch, pink lines the reward zone after the switch.

**Fig. S10**: **Supplementary Figure 3f: Individual cell examples.**

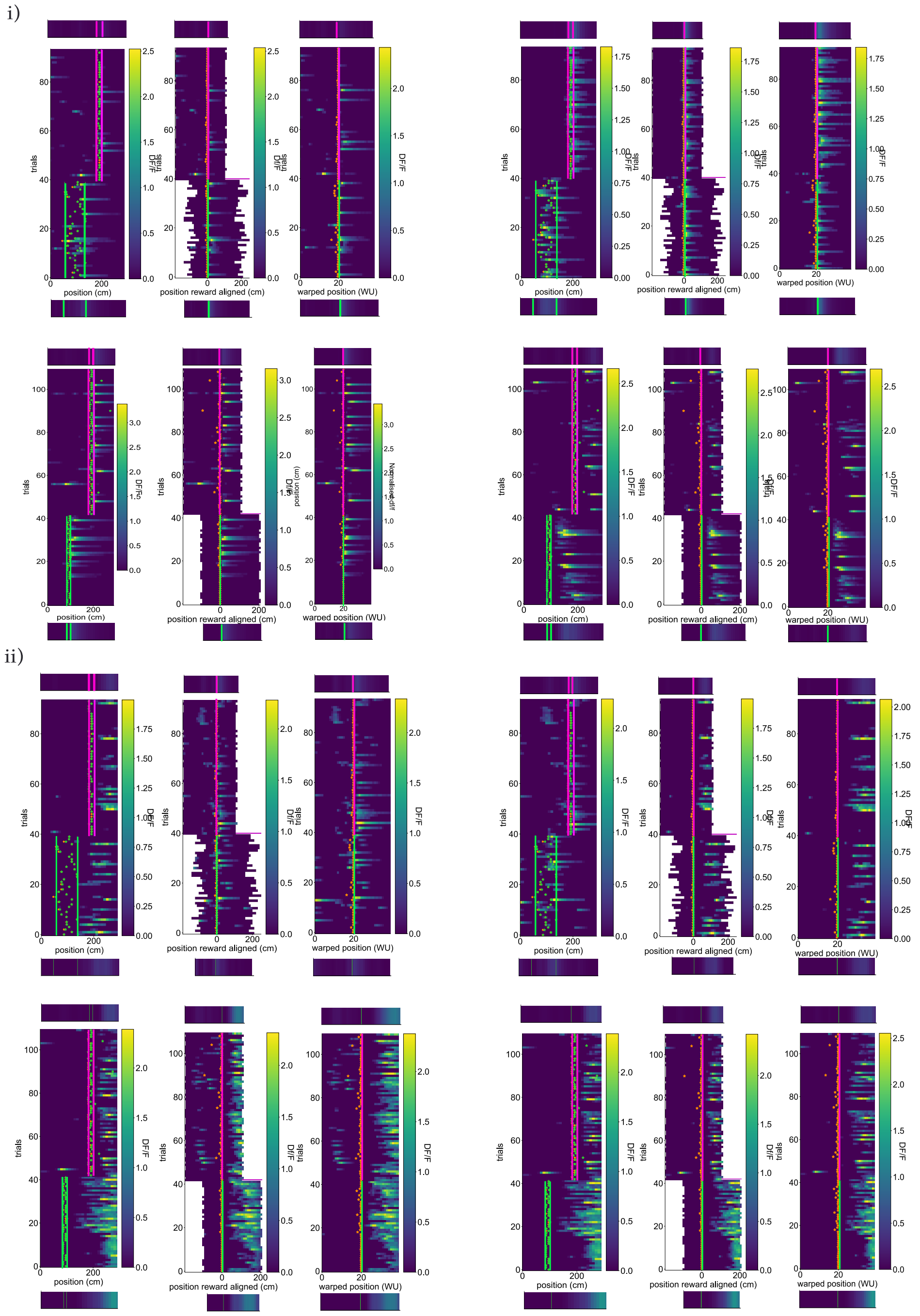

iii)

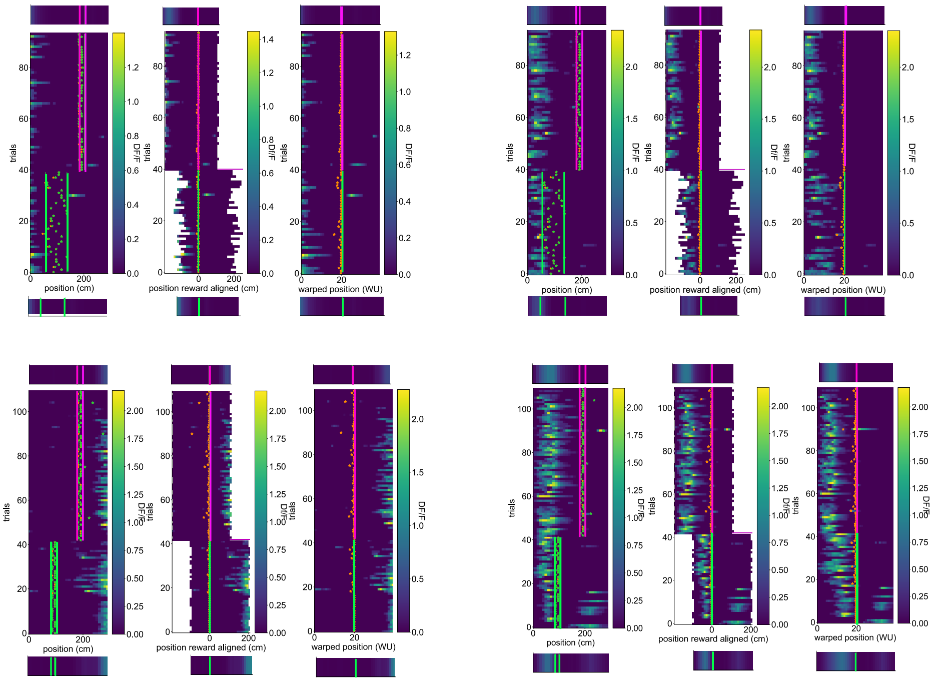


i: Four position-stable cells: example cells that keep their maximum of activity before and after the switch in position reference frame. For each cell, the activity is shown in a position reference frame (left), in a reward reference frame (middle), in a warped reference frame (right). For each plot, the average before and after the switch is shown as insert on top (after) and bottom (before) of the heatmap. Top rows are two cells in UI, bottom rows two cells in UU.
ii: Similar to i: for four reward-stable cells: cells that keep their maximum of activity between before and after the switch in a reward reference frame.
iii: Similar to (i,ii) for four warped-stable cells: cells that keep their maximum of activity between before and after the switch trials in a warped reference frame.
Green thick lines show the full reward zone before the switch, pink lines the reward zone after the switch, green stars the reward consumption locations, and the orange stars the reward delivery location.

---

### Acknowledgements

We are grateful to Claudia Clopath, Matt Jones, Zach Mainen, Tony Pickering and Mark Walton for influential discussions about the design of the task. We thank Marielena Sosa, Mark Plitt and Lisa Giocomo for sharing their results prior to publication.

### Funding

Funding was from the Max Planck Society (CT, PD) and the Humboldt Foundation (PD). PD is a member of the Machine Learning Cluster of Excellence, EXC number 2064/1 – Project number 39072764 and of the Else Kröner Medical Scientist Kolleg “ClinbrAIn: Artificial Intelligence for Clinical Brain Research”. Funding for JRM from Wellcome Trust (101029/Z/13/Z) and Biotechnology and Biological

Sciences Research Council (BBSRC, BB/V001728/1, BB/N013956/1). FX was supported by National Institute of Mental Health Training Program in Neurobiology of Information Storage (T32MH067564) predoctoral fellowship.

### Author Credit Contribution

**Fig. S11**: **CRediT**

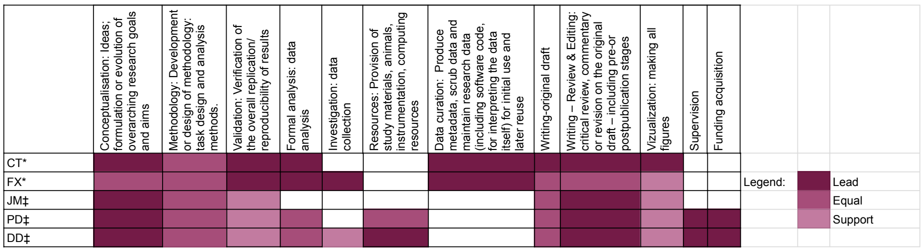


CRediT contribution matrix. Color code refers to the level of contribution per category, as previously used (Tay, 2021). Categories reflect the ones published in the original CRediT taxonomy in (Brand et al, 2015).

### Conflict of interest/Competing interests

The authors declare no conflict of interest.

### References

Anderson MI, Jeffery KJ (2003) Heterogeneous modulation of place cell firing by changes in context. Journal of Neuroscience 23(26):8827–8835

Aoki Y, Igata H, Ikegaya Y, et al (2019) The integration of goal-directed signals onto spatial maps of hippocampal place cells. Cell reports 27(5):1516–1527

Bast T, Wilson IA, Witter MP, et al (2009) From rapid place learning to behavioral performance: a key role for the intermediate hippocampus. PLoS biology 7(4):e1000089

Behrens TE, Woolrich MW, Walton ME, et al (2007) Learning the value of information in an uncertain world. Nature neuroscience 10(9):1214–1221

Best PJ, White AM, Minai A (2001) Spatial processing in the brain: the activity of hippocampal place cells. Annual review of neuroscience 24(1):459–486
